# A Robust Framework for Predicting Mutation Effects on Transcription Factor Binding: Insights from Mutational Signatures in 560 Breast Cancer Genomes

**DOI:** 10.64898/2026.02.14.705876

**Authors:** Hüseyin Hilmi Kılınç, Burçak Otlu

**Affiliations:** Department of Health Informatics, Graduate School Of Informatics, Middle East Technical University, Ankara, 06800, Türkiye

**Keywords:** Breast Cancer, Cancer Genomics, Transcription Factors, Mutational Signatures, Machine Learning

## Abstract

**Background:** A vast majority of somatic mutations in cancer reside in non-coding regions, yet systematically predicting their functional impact on gene regulation remains a significant challenge. These variants often enforce their effects by altering the binding affinity of transcription factors (TFs) to cis-regulatory elements. However, a critical gap exists in linking specific mutational processes to the disruption of gene regulatory networks at a systems level.

**Results:** In this study, we present a comprehensive *in silico* pipeline centered on *k*-mer-based linear regression models to quantify TF binding affinity. Our framework produced 403 high-confidence TF models trained on high-throughput ChIP-seq and PBM datasets. We applied this pipeline to 3.5 million somatic mutations from 560 breast cancer whole genomes to predict gain- or loss-of-function (GOF/LOF) binding perturbations. These predictions were integrated with mutational signature analysis and curated gene sets, utilizing Activity-by-Contact model-based enhancer-gene maps to link variants to their target genes. Our analysis revealed that distinct mutational processes exert non-random, directional effects on specific TF families. The APOBEC-associated signatures (SBS2 and SBS13) were strongly enriched for GOF events in the Myb/SANT and FOX families, while the aging-associated signature SBS1 was enriched for LOF events in the Ets family members. Furthermore, predicted perturbations at putative enhancers were significantly linked to key oncogenes and tumor suppressor genes, with GOF and LOF events (e.g., *FOXA1* and *BRCA1*/*2*, respectively). In breast cancer samples, the basal-like TNBC subtype exhibited that SBS3-driven GOF enrichments for the CXXC family converged on MYC target gene programs, while SBS39-driven LOF events for the same family converged on DNA Repair pathways.

**Conclusions:** Our framework provides a robust and scalable approach for prioritizing and interpreting the functional consequences of somatic mutations in terms of TF perturbations. We demonstrate that specific mutational processes systematically rewire the gene regulatory landscape in a subtype-specific manner, offering novel mechanisms for transcriptional deregulation in breast cancer.

## 1 Introduction

Somatic mutations are alterations in the genomic DNA that arise and accumulate throughout an individual’s lifetime, leaving a molecular record of different endogenous and exogenous mutational processes. These characteristic imprints, particularly those within the vast non-coding regions of the genome, are critical drivers of complex diseases, most notably cancer. While much of cancer genomics has primarily focused on mutations in protein-coding genes, evaluating the oncogenic impacts of non-coding mutations remains a significant challenge [1]. Mutations in these non-coding regions, comprising nearly 98% of the human genome, can potentially perturb the interaction dynamics between transcription factors (TFs) and cis-regulatory elements such as enhancers and promoters, thereby dysregulating gene regulatory networks. Given that the majority of disease-associated variants reside in these non-coding regions, a crucial knowledge gap exists in accurately predicting how they affect TF binding kinetics [2].

In recent years, the application of high-throughput TF-binding assays, such as ChIP-seq [3], HT-SELEX [4], MITOMI [5], and protein binding microarray (PBM) [6], has greatly advanced our ability to characterize transcription factor binding sites (TFBSs). These experimental resources have, in turn, fueled progress in computational prediction of TF binding specificity. Many established methods have relied on Position Weight Matrices (PWMs), which represent TF binding motifs based on nucleotide frequencies [7]. However, PWM-based approaches suffer from several drawbacks that limit their predictive power: they are prone to high false-positive rates, often identifying non-functional genomic sequences as potential binding sites, and they also operate on the simplifying assumption that nucleotides within a binding site contribute independently to the overall binding affinity [8–10]. Therefore, more recent studies have shifted beyond these motif-based models to (i) conventional machine learning approaches [11–13], (ii) deep learning frameworks such as DeepBind [14], FactorNet [15], and (iii) transformer-based models such as BERT-TFBS [16], and EpiGePT [17], which better capture the complexity of TF–DNA interactions.

At the same time, mechanistically and clinically, understanding the functional impact of somatic mutations on TF binding must also be viewed through the lens of mutational processes. Mutational signature analysis has emerged as a powerful framework for dissecting the endogenous and exogenous processes that shape the instability of cancer genomes. Each mutational signature is a characteristic footprint of DNA damage that different mutational processes engrave across cancer genomes for single-base substitutions (SBS), doublet-base substitutions (DBS), small insertions and deletions (indels), copy-number changes, and structural rearrangements [18]. Integrating mutational signatures with TF-binding perturbations confers a principled route to connect distinct mutational processes to regulatory dysfunction. Previous studies often focused either on identifying recurrent mutations at regulatory hotspots or on predicting TF-binding changes in isolation [19, 20]. However, despite these advances, a systematic integration of TF-binding perturbations with mutational signatures and the regulatory map is still less explored. Besides this, breast cancer (BC) is known as one of the most diagnosed malignancies worldwide and a molecularly heterogeneous disease [21]. The contribution of TF perturbations to molecular subtype-specific regulatory mechanisms in breast cancer remains poorly understood. Accordingly, the molecular heterogeneity of breast cancer makes it a particularly compelling system in which to study subtype-specific regulatory disruptions driven by perturbed TF dynamics.

In this study, we address this gap by developing a machine learning-based pipeline that quantitatively predicts gain- or loss-of-function (GOF/LOF) events for 403 distinct human TF across somatic mutations. The pipeline leverages *k* -mer-based linear regression models trained on both *in vivo* (ChIP-seq) and *in vitro* PBM datasets and employs the stochastic gradient descent (SGD) optimization to estimate coefficients of model features. We applied this framework to 560 breast cancer samples, a landmark cohort previously published by Nik-Zainal and colleagues [22]. For any given variant, the trained models compute the predicted change in TF binding affinity and assign calibrated statistical significance. To connect distal regulatory changes to target genes, we leverage enhancer–gene maps based on the Activity-by-Contact (ABC) model, enabling assignment of predicted TF-binding perturbations to putative target genes [23]. To place these predictions in a biological context, we systematically integrated them into breast cancer molecular subtypes. In this study, we aim to uncover how distinct mutational processes preferentially perturb specific TF families and converge on critical oncogenic and tumor suppressive pathways. By connecting mutational signatures to TF binding alterations and downstream enhancer–gene programs, this study provides a systematic map of regulatory perturbations in breast cancer. Furthermore, we uncover associations between specific mutational signatures and TF families, highlight potential regulatory drivers, and reveal distinct subtype-specific patterns of disruption. Taken together, our work contributes to the growing effort to understand the regulatory consequences of non-coding mutations in cancer and establishes a framework for connecting mutational signatures to gene regulatory dysfunction.

## 2 Methods

### 2.1 Datasets

Predictive models of TF binding specificities were trained on a comprehensive set of high-throughput experimental datasets, covering both *in vivo* and *in vitro* modalities. *In vivo* dataset from TF ChIP-seq experiments was retrieved from the ENCODE Project database (released/approved status only) [24]. We specifically retained experiments with bed narrowPeak formatted files, which report genomic binding peaks along with their corresponding enrichment scores (signalValues). A total of 2557 TF-ChIP-seq datasets across diverse human cell lines and tissues were collected after filtering for high-quality status and excluding experiments with severe “audit” flags. We also used *in vitro* dataset from universal protein-binding microarray (uPBM) assays that measure fluorescence intensity of a specific TF bound to a comprehensive library of double-stranded DNA probes (a total of ≈ 40,000). These experiments were designed with 60-nt de Bruijn sequences covering all possible 10-mers [6]. In total, 588 PBM data for various human TFs were sourced from UniPROBE and CIS-BP databases [25, 26].

### 2.2 Sequence-Based TF Binding Models

#### 2.2.1 Feature Construction and *k* -mer Selection

This study is built on a *k* -mer-based linear regression model to predict TF binding specificity, where each DNA sequence is represented by frequencies of overlapping k-mers. In this model, *k* -mers are used as features to represent the sequence-dependent determinants of TF binding. Selecting the most suitable *k* value is a critical step, as it directly influences the accuracy and reliability of predictive TF-binding models. Given the four nucleotide bases and all possible *k* -mer combinations, with reverse-complement collapsing to account for the strand-agnostic nature of TF binding to double-stranded DNA, the total number of unique features was defined as follows:

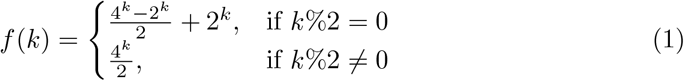

While choosing the optimal *k* value for our models, we considered two primary factors: (i) motif-level biological relevance, requiring *k* to be sufficiently long to capture the specificity of most TF motifs (typically ≈ 6-12 bp), and thus excluding shorter *k* - mers (*k* ≤ 5) that oversimplify sequence context and lead to underfitting; (ii) tractable feature dimensionality, where model size and learnability are governed by the number of *k* -mer features, motivating the exclusion of larger *k* -mers (*k* ≥ 7) due to the exponential growth in dimensionality, which increases overfitting risk when (*p* ≥ *n*), exacerbates the curse of dimensionality, and substantially inflates computational and memory costs. Balancing these considerations, we selected (*k* = 6) as an operating point that preserves biological signal and results in 2,080 reverse-complement-collapsed features while maintaining a manageable feature space for robust model training. For each input sequence (e.g., 60-nt PBM probe or ChIP-seq peak window), a sliding window enumerated all overlapping 6-mers (max 55 per 60-nt sequence). We constructed per-sequence frequency vectors (2,080-dimensional), applied min–max scaling to input features, and used log2-transformed binding signal intensities as targets.

#### 2.2.2 Model Training and Uncertainty Estimation

The relationship between 6-mer frequencies and TF binding affinity was modeled using a multiple linear regression framework:

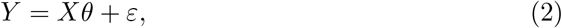

where *X* is the scaled 6-mer frequency matrix (occurrence counts), *θ* is the coefficient vector for 6-mer binding contributions, *Y* represents observed signal intensities, and *ε* denotes the error term.

We employed the Stochastic Gradient Descent (SGD) method to train the regression models due to its computational efficiency in handling the large and sparse datasets. The SGDRegressor module from the Python scikit-learn library was used with tuned hyperparameters (*learning rate* (*η*) = 0.1, *max iter* = 1000, *tol* = 0.001, *alpha* = 0.0001), optimizing mean squared error during training. After estimating the optimal coefficients *θ* via SGD, the coefficient covariance matrix and the model’s residual variance (*σ*^2^) were computed as:

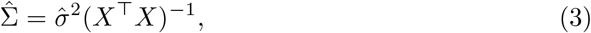

where residual variance 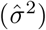 is defined as 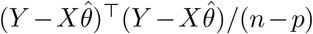 with *n* being the number of observations (sequences), which varies depending on the TF datasets, and *p* the number of 6-mer features (*p* = 2,080). The variance 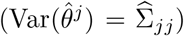 and the standard error 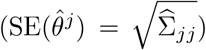 are obtained from the diagonal of this *p × p* matrix. This formulation confers the propagation of model uncertainty to downstream analyses, allowing the statistical significance of predicted binding-affinity changes to be scaled by the propagated variance derived from 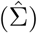.

### 2.3 Model Evaluation and Selection

We initially compiled a total of 3,145 TF-specific datasets (2,557 ChIP-seq and 588 PBM). Datasets with fewer observations than features (265 ChIP-seq datasets with *<* 2,080 peaks) were excluded. Individual regression models were trained for each remaining TF dataset. Model performance was assessed using 10-fold cross-validation, with (*R*^2^) as the performance metric. To ensure model quality, we applied a mean threshold of *R*^2^ *>* 0.1 for models trained on TF ChIP-seq data and a more stringent threshold of *R*^2^ *>* 0.15 for models trained on PBM data. A total of 436 models (187 from ChIP-seq-based, 249 from PBM-based) surpassed these thresholds. For the same type of TFs with multiple qualifying models, the model with the highest *R*^2^ score was selected. This rigorous selection process yielded a final set of 403 unique, high-confidence human TF models, whose estimated coefficients 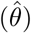 and their covariance matrices 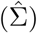 were stored for downstream analyses.

### 2.4 Predicting Mutation Effects on TF Binding

#### 2.4.1 Quantifying Binding Affinity Change

The pre-trained curated models were used to predict the impact of any given somatic mutation on TF binding affinity. In this study, we focused on single-base substitutions (SBS), where a mutation can perturb overlapping 6-mers within an 11-bp window centered on the mutation site. The change in TF–DNA binding affinity (Δ*B*) was calculated as a weighted linear combination of the changes in overlapping 6-mer frequencies between the mutated and wild-type sequences:

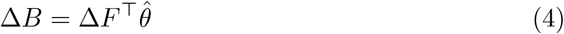

Here, Δ*F* denotes the contrast vector representing the change in 6-mer frequencies (*F*_mutated_ − *F*_wild-type_) within the 11-bp window context, weighted by the estimated coefficients 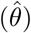. The statistical significance of each predicted binding change was assessed using a t-test under the null hypothesis (*H*_0_ : Δ*B* = 0). For this test, we assumed the model’s error term follows a normal distribution: *ϵ* ∼ *𝒩*(0, *σ*^2^*I*), and the t-statistic was calculated as:

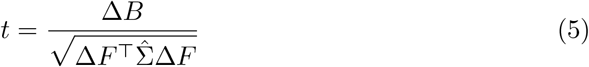

This statistical framework inherently accounts for each model’s uncertainty, ensuring that weaker models (bearing higher residual variances and thus larger entries) require a larger Δ*B* score to achieve statistical significance. Given the large number of observations (*n* ≫ *p*) in our training data, the t-statistic approximates a z-score, which was then used to derive a p-value for each predicted binding change. Mutations exhibiting a p-value ≤ 0.01 corrected via the Bonferroni correction were considered statistically significant and were subsequently classified as gain-of-function (GOF) or loss-of-function (LOF) events based on the direction of the Δ*B* score. As a result, for a mutation to be deemed statistically significant in such models, it must induce a more substantial change in binding affinity (Δ*B*). This self-correcting feature ensures that the final predictions remain robust and properly control for model-specific uncertainty.

### 2.5 Analysis of the Breast Cancer Cohort

#### 2.5.1 Somatic Mutation Profiling and Mutational Signature Extraction

We applied our pipeline to a dataset of 3,479,652 single-base substitutions identified from whole-genome sequencing of 560 breast cancer samples [22]. Mutational processes were characterized using the SigProfilerExtractor tool [27] to decompose into COSMIC (Catalogue of Somatic Mutations in Cancer, v3.4 database [28]) mutational signatures. This analysis aims to identify the characteristic footprints left by different mutational processes, such as exposure to exogenous mutagens or defects in endogenous biological pathways. This process extracted the mutational signatures and assigned a probability for each signature’s contribution to individual mutations.

#### 2.5.2 Simulated Null Mutation Landscapes for Enrichment Assessment

To assess the significance of our downstream enrichment results, we incorporated SigProfilerSimulator [29], a complementary tool within the SigProfiler suite that generates biologically realistic null mutation landscapes. For each breast cancer sample, the complete somatic mutation catalog was simulated *n* times (100 iterations by default), redistributing mutations across the genome while preserving key properties of the original mutation spectrum, including chromosome-level mutation counts, total mutational burden, and profile (e.g., trinucleotide context) at the specified resolution. Both observed and simulated mutations were subsequently classified into mutation types and probabilistically assigned to mutational signatures using identical procedures. By integrating SigProfilerSimulator, we established an empirical null distribution to model background mutation patterns. Comparing observed TF binding perturbations against this baseline allowed us to identify significant enrichments or depletions beyond mutational noise.

#### 2.5.3 Linking Regulatory Mutations to Target Genes

To link mutations in distal regulatory elements to their target genes, we utilized pre-computed enhancer-gene maps from the Activity-by-Contact (ABC) model [23]. We leveraged predicted enhancer-gene connections combined from relevant breast cancer cell lines and biosamples (e.g., MCF-7, MCF-10A, MDA-MB-231). Significant TF binding perturbations were first mapped to putative enhancer regions within these cell lines. The ABC model predictions were then used to link these enhancers to their target genes, as schemed in Fig. 1. Finally, we cross-referenced these target genes with the OncoKB database [30] to identify regulatory perturbations affecting known oncogenes and tumor suppressor genes.

**Fig. 1.**
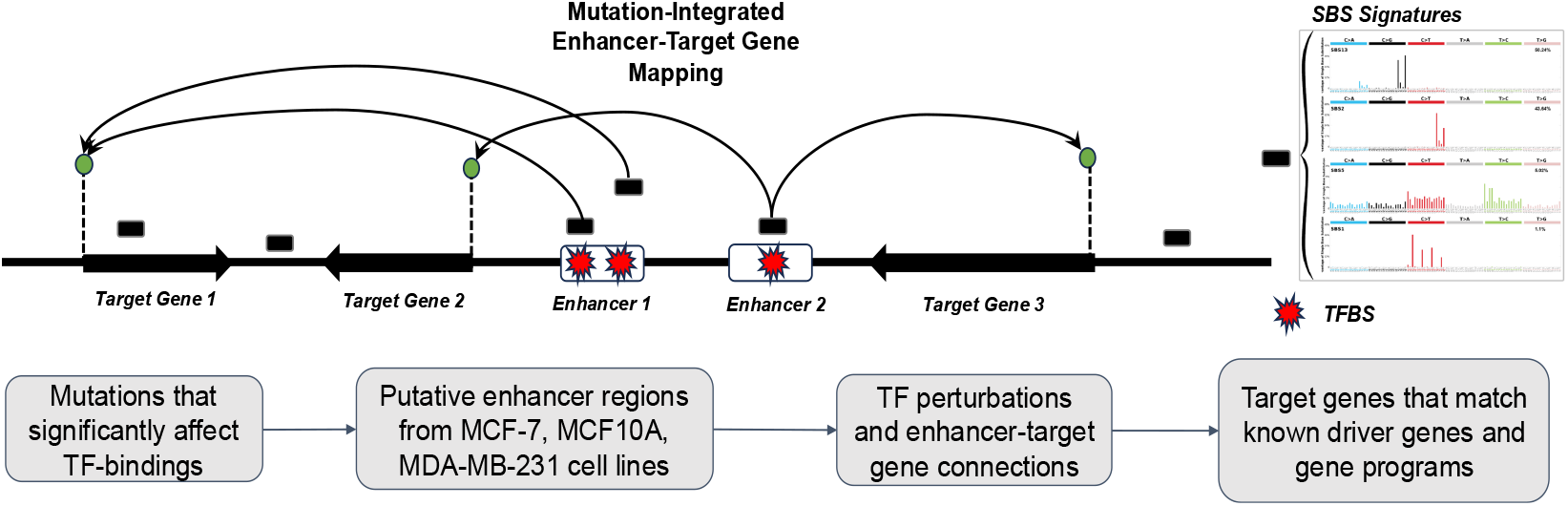
Schematic overview of the mutation-integrated enhancer–target gene mapping via the Activity-by-Contact model.

## 3 Results

### 3.1 Performance of Predictive TF Models

The final set of 403 predictive TF models achieved a mean *R*^2^ of 0.39 (median = 0.35) across 10-fold cross-validation (Fig. S1). As expected, models trained on well-designed *in vitro* PBM data generally achieved higher *R*^2^ scores than those trained on noisier *in vivo* ChIP-seq data, which is reflected in our tiered selection thresholds (*R*^2^ *>* 0.15 for PBM, *R*^2^ *>* 0.1 for ChIP-seq). This strategy balanced the need for high-quality models to achieve broad coverage across the human TF repertoire. Further analysis (Fig. 2) revealed that model performance varied considerably across different TF families, which are groups of TFs sharing structurally similar DNA-binding domains (DBDs). Certain families demonstrated high predictability. For instance, the TATA-box binding protein (TBP) family exhibited the highest mean *R*^2^ score at 0.7, reflecting exceptionally strong model performance. Other large and well-characterized families, including Homeodomain, Ets, and CxxC, also yielded models with high mean *R*^2^ values, indicating that the binding specificities of their members are well-captured by the *k* -mer-based model with SGD optimization. Conversely, families such as TEA, MYM-type ZF, and SMAD displayed lower mean *R*^2^ scores, suggesting that their DNA-binding mechanisms may be more complex or less dependent on local 6-mer sequence context, posing a greater challenge for this modeling framework.

**Fig. 2.**
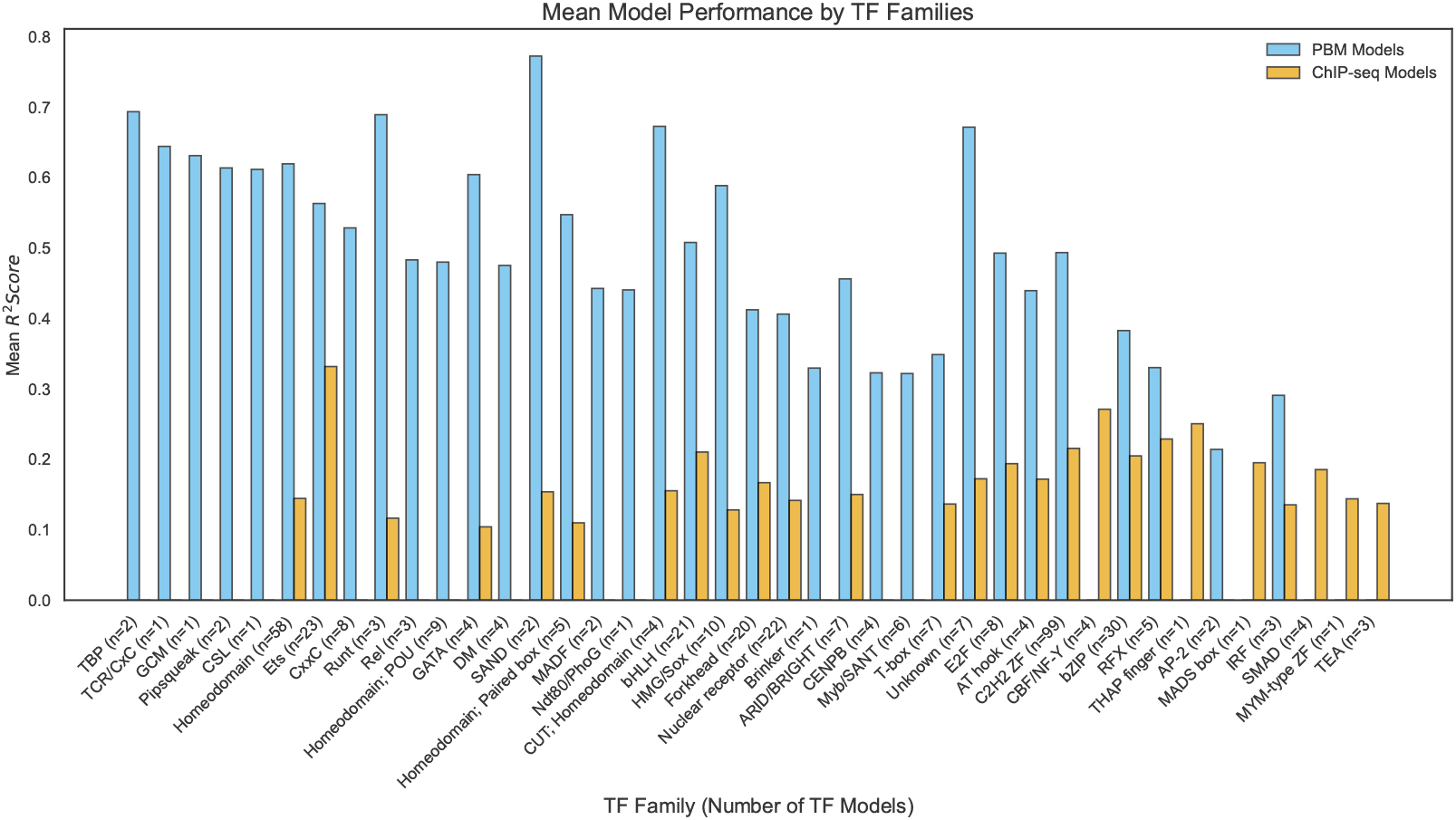
Average *R*^2^ scores of ChIP-seq-based and PBM-based models grouped by TF DBD family.

### 3.2 Landscape of TF Binding Perturbations in Breast Cancer Cohort

To systematically quantify the functional impact of somatic mutations on TF-DNA interactions in a cancer context, the validated set of 403 TF binding models was applied on a comprehensive dataset of ≈ 3.5 million single-base substitutions identified from the whole-genome sequences of 560 breast cancer samples. For each mutation’s effect on each of the TF dynamics, a binding perturbation score (Δ*B*) was measured. To illustrate the biological interpretation of these quantitative predictions, two representative examples from the breast cancer cohort are presented in Fig. 3. The first case demonstrates a significant GOF event (Fig. 3A). A single-base substitution (A*>*C) in sample PD4976a was predicted to substantially increase the binding affinity for the TF MYBL2. This prediction is supported by the mutation’s effect on the local sequence, which creates a *de novo* stronger match to the consensus MYBL2 binding motif (6.93 Δ*B* score, adjusted *p* value ≤ 0.01). The second example showcases a significant LOF event (Fig. 3B). C*>*T mutation in sample PD7322a was quantitatively predicted to disrupt an existing binding site for the TF ETS1, resulting in a significant decrease in its binding affinity (-7.52 Δ*B* score, adjusted *p* value ≤ 0.01).

**Fig. 3.**
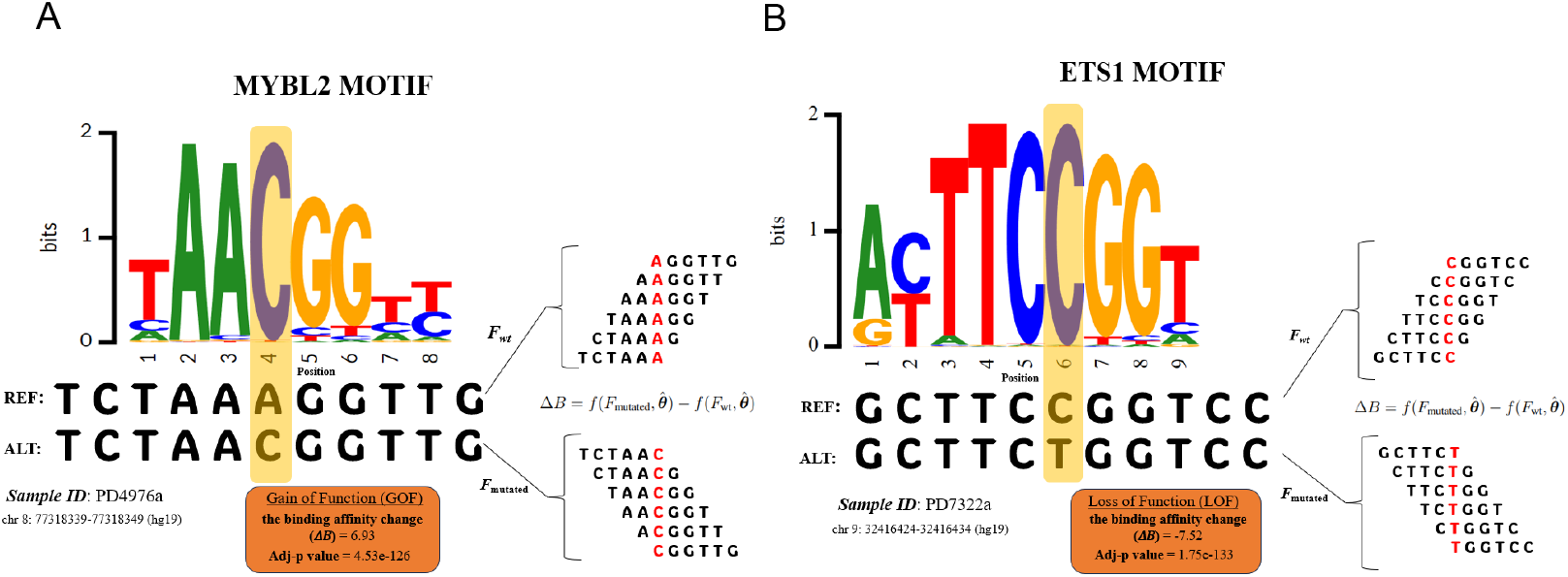
Single-base substitutions from breast cancer samples (PD4976a and PD7322a) matched the reference motifs from Factorbook [31]. **A** Gain-of-binding due to an A*>*C mutation overlapping the MYBL2 motif, showing a statistically significant increase in predicted binding affinity. **B** Loss-of-binding caused by a C*>*T mutation, significantly reducing ETS1 affinity.

In both instances, the predicted functional consequences align with visual inspection of the reference and mutated sequences against the canonical TF motif examples curated in the Factorbook database [31], providing qualitative support for the model’s quantitative predictions. These examples highlight the model’s capacity to pinpoint specific mutations that likely alter the gene regulatory landscape by modifying distal TF binding sites.

### 3.3 Assignment of Operative Mutational Signatures in Breast Cancer Samples

#### 3.3.1 *De novo* Extraction and Decomposition

To characterize the aetiological drivers underlying regulatory changes, a mutational signature analysis was performed on the cohort of 560 breast cancer genomes. Utilizing the SigProfilerExtractor tool, we performed *de novo* extraction of single-base substitution (SBS) signature and identified 12 operative SBS signatures as the optimal solution based on reconstruction accuracy and signature stability (selection plot in Fig. S2). These 12 extracted *de novo* signatures were then decomposed into COSMIC v3.4 reference signatures to enable attribution to known mutational processes.

The decomposition mapped to 16 COSMIC SBS signatures, which were active at least one or more samples. After further filtering out signatures with low-confidence assignments present in very few samples (SBS17a and SBS21), a final set of 14 SBS signatures was retained for downstream analyses.

The prevalence and relative contribution of these 16 COSMIC SBS signatures across the cohort are summarized using a tumor mutational burden (TMB) plot (Fig 4). TMB analysis revealed that signatures SBS1, SBS2, SBS5, and SBS13 were ubiquitously active across the BC cohort, whereas the influence of other signatures was localized to specific cases, showing a more sporadic distribution.

**Fig. 4.**
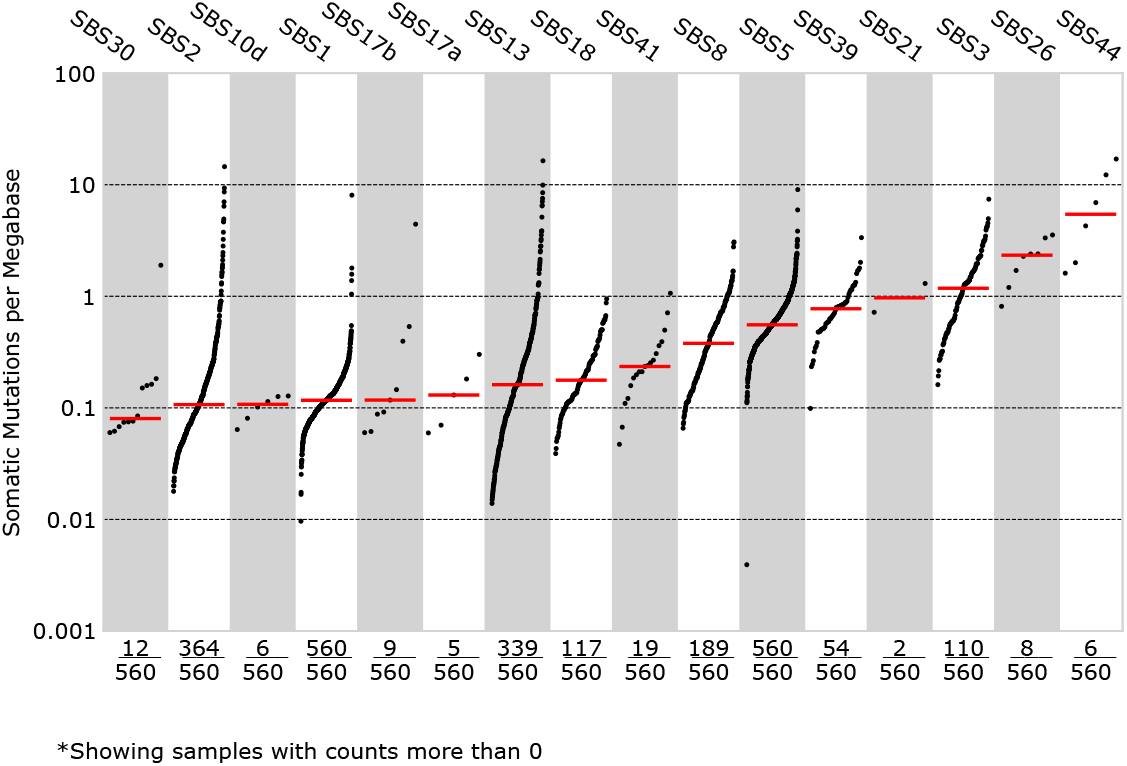
Tumor Mutational Burden (TMB) plot illustrating the contribution of each mutational signature across samples.

#### 3.2.2 Mutational Signature Activities Across Molecular Subtypes of Breast Cancer

To examine whether the activity of mutational signatures varies across breast cancer molecular subtypes, samples from the cohort were stratified into Luminal A, Luminal B, HER2-enriched, and Triple-Negative Breast Cancer (TNBC). Luminal A represented the largest fraction of samples (57.5%), followed by TNBC (29.9%), as shown in Fig. 5. Analysis of signature activity within these subtypes revealed distinct patterns. Signature SBS5, attributed to a clock-like signature, was found to be the most prevalent signature across all samples, with slightly higher activity in Luminal A tumors. The APOBEC3-associated signatures SBS2 and SBS13 showed widespread activity across the cohort, with a marked prevalence in the HER2-enriched subtype compared to other groups (Fig. 5). A particularly striking finding was the significant enrichment of SBS3 in the TNBC subtype. SBS3 is the characteristic signature of homologous recombination deficiency (HRD), commonly associated with *BRCA1/2* dysfunction [22]. Its prevalence within TNBC supports a well-established hallmark of this aggressive subtype [32–34]. The clear emergence of this pattern in our cohort supports the biological plausibility of the extracted signatures and indicates that SBS3-driven mutations are likely to play a central role in subtype-specific regulatory perturbations in the pathogenesis of TNBC.

**Fig. 5.**
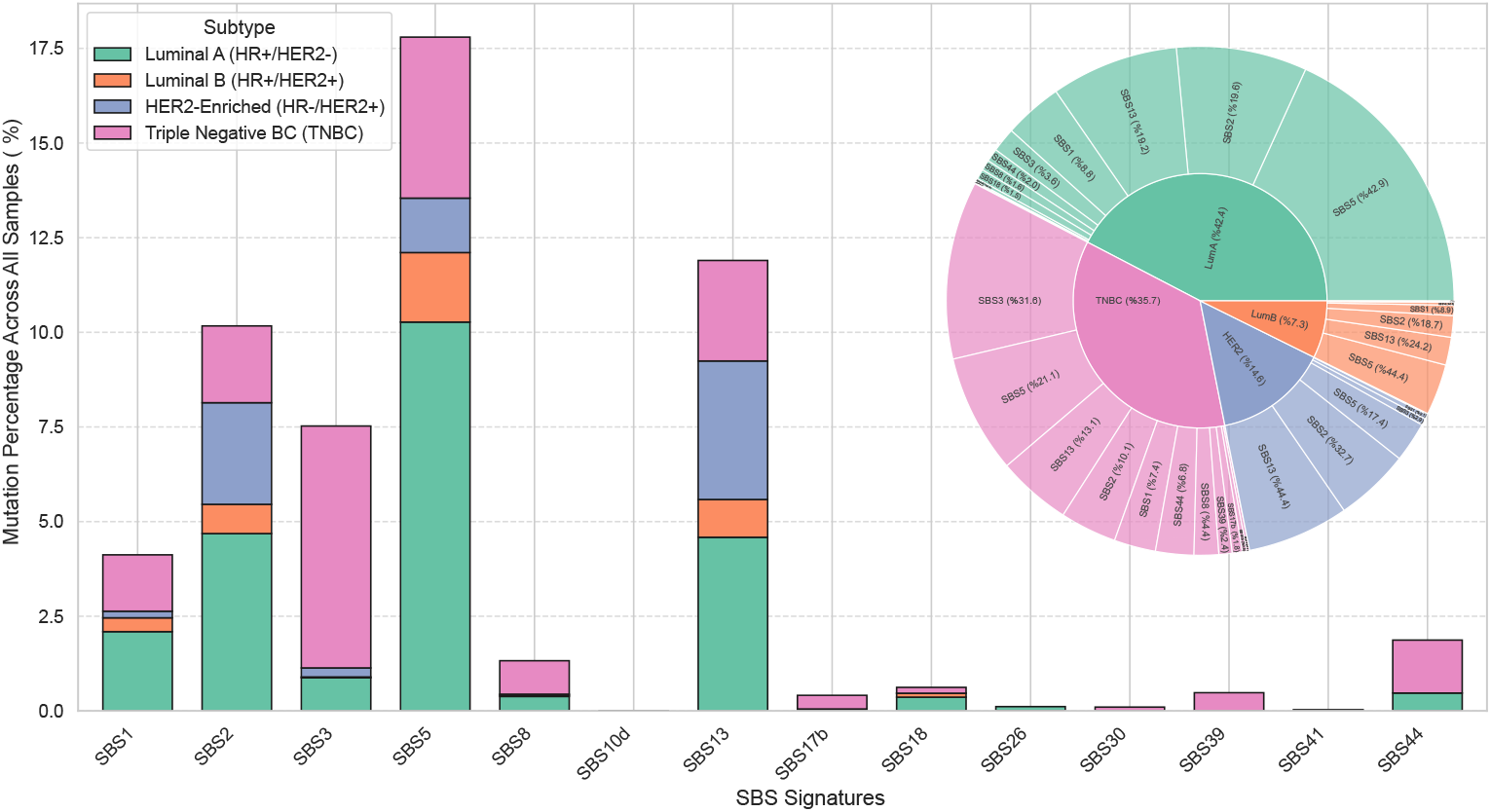
Distribution and composition of SBS signatures across breast cancer subtypes in 560 samples. Percentages represent the proportion of mutations attributed to each signature. The inset sunburst plot summarizes the overall compositional distribution of SBS signatures within each molecular subtype.

#### 3.3.3 Systematic Association of Signatures with TF Binding Perturbations

We probabilistically attributed each of the ≈ 3.5 million somatic mutations to a single best-fit signature, setting a minimum signature probability threshold of 0.75 to ensure high-confidence attribution. A summary of the 14 active SBS signatures, including their proposed aetiologies, the total number of high-confidence mutations attributed to each, and the numbers of GOF, LOF, and neutral effects predicted from the aggregated behavior of all 403 TFs, is provided in Table 1. To statistically test for associations between mutational signatures and overall binding directions of TFs, Fisher’s exact test was conducted to compare the observed GOF-to-LOF ratios for each TF–signature pair against the null expectation of no association. For all subsequent analyses, an association was deemed significant if it met the criteria of a Bonferroni-corrected *p* value ≤ 0.05 and an absolute log2FC (gain/loss) ≥ 0.25. TFs not meeting these criteria were classified as neutral.

**Table 1.** Summary of mutational signatures and their transcription factor perturbation statistics.

| Signature | Etiology / Process | Mutations | TFs | GOF | LOF | Neutral |
| --- | --- | --- | --- | --- | --- | --- |
| SBS1 | Spontaneous deamination of 5-methylcytosine | 146118 | 403 | 180 | 117 | 106 |
| SBS2 | APOBEC activity | 361798 | 403 | 186 | 114 | 103 |
| SBS3 | HR deficiency | 262069 | 403 | 149 | 44 | 210 |
| SBS5 | (Unknown) Aging/Tobacco smoking/NER deficiency | 633021 | 403 | 117 | 102 | 184 |
| SBS8 | (Unknown) HR/NER deficiency | 47897 | 403 | 194 | 55 | 154 |
| SBS10d | Defective POLD1 proofreading | 177 | 403 | 77 | 1 | 325 |
| SBS13 | APOBEC activity | 423962 | 403 | 206 | 67 | 130 |
| SBS17b | Damage by ROS/5FU chemotherapy | 14314 | 403 | 98 | 123 | 182 |
| SBS18 | Damage by ROS | 22429 | 403 | 184 | 45 | 174 |
| SBS26 | MMR deficiency | 3921 | 403 | 49 | 129 | 225 |
| SBS30 | BER deficiency | 3488 | 403 | 115 | 45 | 243 |
| SBS39 | Unknown | 17622 | 403 | 57 | 60 | 286 |
| SBS41 | Unknown | 924 | 403 | 4 | 6 | 393 |
| SBS44 | MMR deficiency | 65099 | 403 | 222 | 45 | 136 |

A total of 2,002,839 mutations were assigned to SBS signatures with probability $> 0.75$ .

### 3.4 Signature-Specific Trends in TF Family-Level Perturbations

Analyzing perturbations at the TF family level captures broader, coordinated patterns of regulatory disruption, grouping TFs by shared DNA-binding domain (DBD) architecture. This aggregated approach allows for a systematic assessment of whether specific mutational signatures drive biased binding gains or losses across entire TF families. A global overview of family-level effects is shown in Fig. S3 using a heatmap of pie charts summarizing the aggregate impact of each of the 14 mutational signatures on individual TF families across all breast cancer samples. For each signature–TF family pair, pie chart segments quantify the proportion of TF family members exhibiting significant gain-of-function (red), loss-of-function (blue), or non-significant (gray) binding effects. This analysis revealed pronounced non-random patterns, with several mutational signatures displaying strong directional biases toward either GOF or LOF perturbations in specific TF families.

#### 3.4.1 Preferential Effects of APOBEC Signatures

One of the most prominent patterns to emerge from the family-level perturbations was the specific impact of the APOBEC-associated signatures (SBS2 and SBS13) on certain TF families. Aberrant activity of the APOBEC3 enzymes induces cytosine deamination, leading to C*>*T and C*>*G mutations within a TpC sequence context, which are especially accumulated in breast cancer genomes [35]. The heatmap of pie charts (Fig. 6A) illustrates a pervasive gain-of-function (GOF) trend, indicated by the predominantly red coloring, for Forkhead box (FOX) family TFs specifically under the common influence of SBS2 and SBS13. Volcano plots for SBS2 and SBS13 (Fig. 6B, C) reveal a statistically robust trend toward gain-of-function enrichment within the FOX family with high statistical significance (*q* ≪ 0.05, |log_2_ FC| ≥ 0.25). These findings demonstrate that a significant enrichment of APOBEC-induced mutations creates *de novo* FOX family binding sites relative to matched background expectations.

**Fig. 6.**
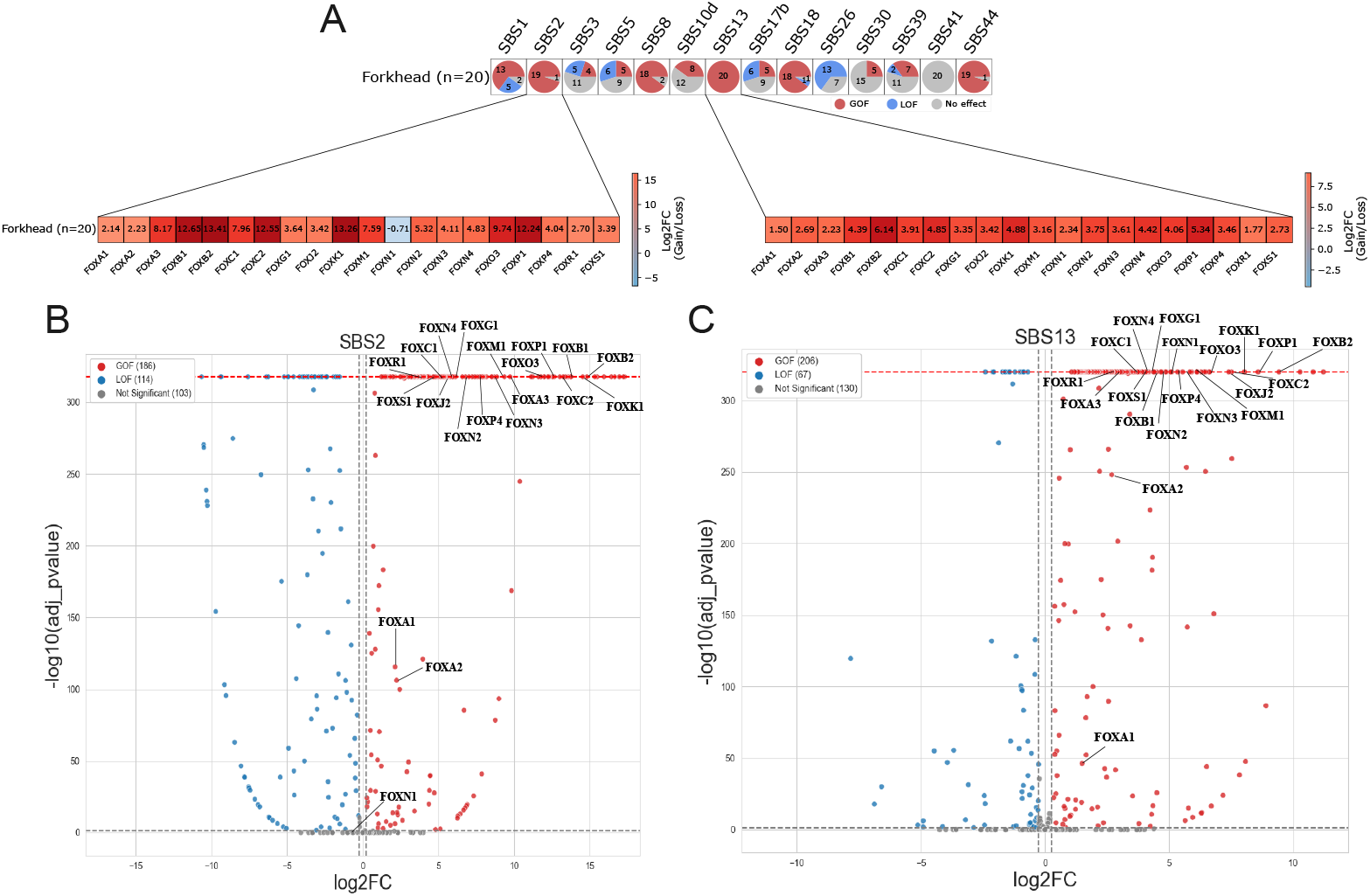
The enrichment of differentially perturbed Forkhead box (FOX) family members by APOBEC signatures. **A** The heatmap of pie charts displaying the proportion of individual FOX family members across multiple mutational signatures (GOF = red, LOF = blue, non-significant = gray). **B** Volcano plot showing SBS2-associated perturbations with labels displayed exclusively for FOX family members. **C** Volcano plot for SBS13-associated perturbations, following a similar format as panel (B) and highlighting a similar GOF enrichment for members of the Forkhead family.

Beyond the FOX family TFs, a similar strong GOF enrichment was significantly observed for the Myb/SANT family members in response to APOBEC-driven mutations. Volcano plots for SBS2 and SBS13 (provided in Fig. S4) show a marked shift of Myb/SANT family members toward GOF effects, suggesting that APOBEC-associated substitutions preferentially enhance MYB-like binding motifs. This observation is consistent with prior reports linking APOBEC activity to increased binding affinity of MYB and MYBL2 family members, driving abnormal activation of nearby oncogenes [36–38]. This also supports the sensitivity of our framework to detect signature-specific regulatory perturbations.

#### 3.4.2 Preferential Effects of SBS3 Signature

Another distinct perturbation pattern emerged for the SBS3 signature, which has been previously associated with HRD and is commonly enriched in *BRCA1/2* -mutated breast cancers [22]. SBS3-driven mutations produced a clear family-level enrichment of GOF events, predominantly affecting the CxxC domain–containing TFs. All eight members of this family exhibited significantly positive log_2_ fold changes, indicating enhanced predicted binding activity in the presence of SBS3-associated mutations (Fig. 7A, B). The zoomed heatmap view in Fig. 7A reveals a strong GOF bias across the CxxC family, while the corresponding volcano plot in Fig. 7B shows consistent clustering of CxxC TFs in the GOF-enriched region with high statistical significance (*q <* 0.05, |log_2_ FC| ≥ 0.25).

**Fig. 7.**
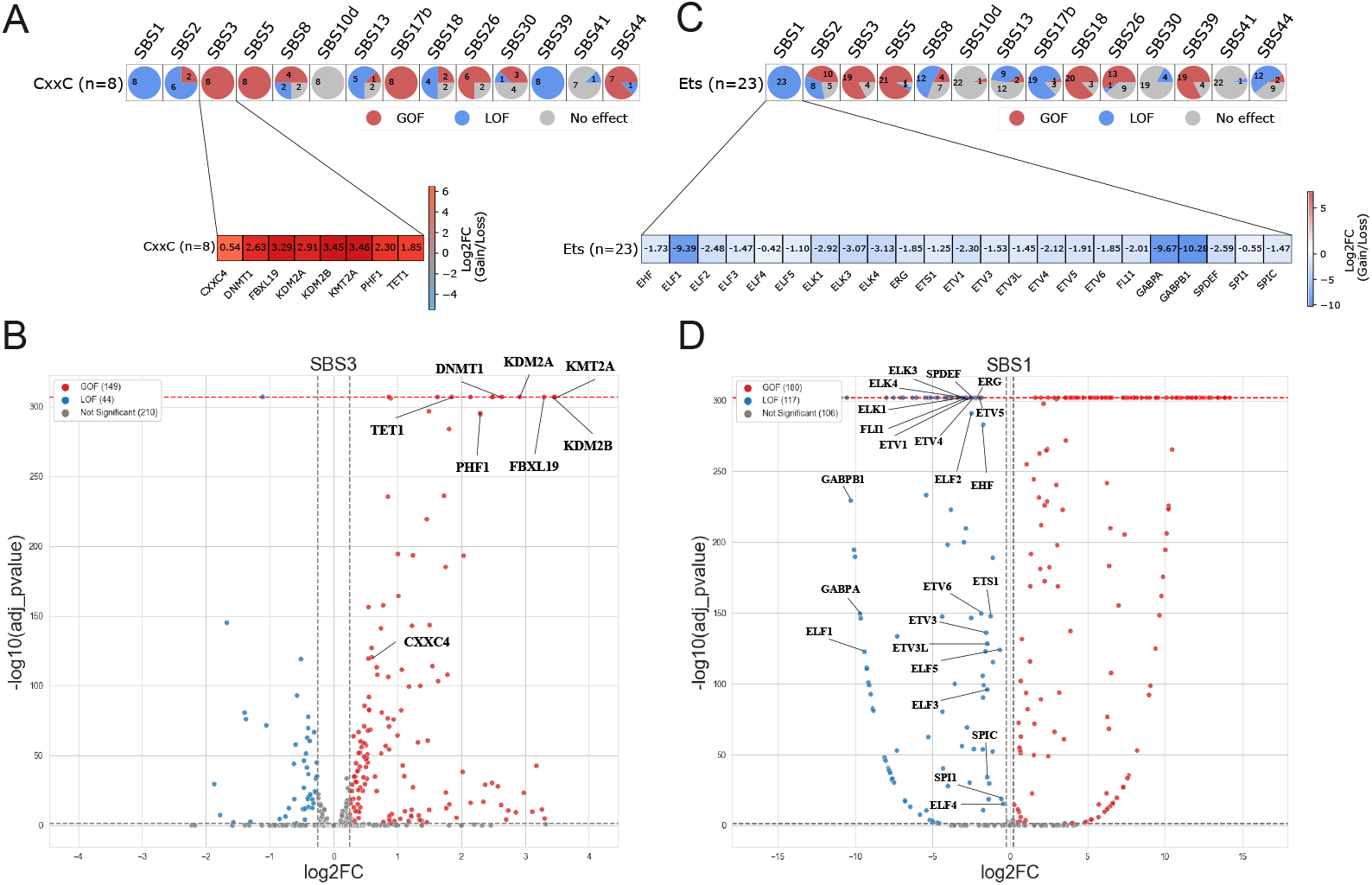
Enrichment of differentially perturbed Ets and CxxC family members due to SBS1 and SBS3 mutations. **A** Heatmap of CxxC TFs under SBS3, revealing a predominant GOF pattern. **B** Volcano plot for SBS3-associated perturbations, demonstrating significant GOF enrichment for CxxC TFs. **C** Heatmap showing predominant LOF pattern across Ets TFs, indicating a dominant LOF trend for SBS1. **D** Volcano plot for SBS1-associated perturbations, highlighting significant LOF events among Ets members.

It is worth noting that the companion HRD-linked signature, SBS39, displayed a reverse directional effect on the same TF family, with most CxxC members showing LOF perturbations in the previous aggregated pie-chart analysis (see Fig. S3). Rather than being contradictory, this functional divergence may reflect a partitioned role in tumor evolution, wherein each signature disrupts independent biological modules necessary for the phenotype.

#### 3.4.3 Preferential Effects of SBS1 Signature

In contrast to the GOF-dominant patterns observed for other signatures, our analysis for the SBS1 signature identified a significant and inverse trend specifically across the Ets family TFs. SBS1 is a clock-like signature resulting from the spontaneous deamination of 5-methylcytosine, a mutational process that accumulates gradually with age and predominantly affects CpG dinucleotides [39]. Our signature-specific results in the family-level aggregation revealed that mutations attributed to SBS1 consistently weaken Ets binding affinity. In the heatmap of pie charts (Fig. 7C), all Ets family members exhibited negative log_2_ fold changes, indicating a uniform LOF bias. This trend can be also seen in the corresponding volcano plot (Fig. 7D), in which all Ets members clustered toward the LOF-enriched region with statistical significance (*q <* 0.05, |log_2_ FC| ≥ 0.25). Collectively, these all SBS-specific findings align with the broader concept that mutational processes not only drive genetic alterations but also impose directional pressures on regulatory DNA by selectively altering TF-binding affinities in the family context.

### 3.5 Integration of Operative Mutations with Enhancer-Target Gene Maps

To translate predicted binding changes into potential effects on gene regulation, we integrated operative mutations with genome-wide enhancer-gene maps derived from the Activity-by-Contact (ABC) model predictions [23]. This facilitated the mapping of functional, TF-perturbing mutations within putative enhancers to their respective ABC-predicted target genes. In total, 45,324 breast cancer mutations satisfied both criteria of (i) overlapping ABC-defined enhancers active in breast cancer–relevant cell lines and (ii) inducing statistically significant TF binding gain or loss (adjusted p ≤ 0.01, per our earlier predictions). These mutations mapped to 164,290 mutation–target gene pairs, spanning a broad range of housekeeping and tissue-specific genes. To investigate the most clinically critical cancer-related events in this pipeline, these enhancer mutations were further filtered to retain only those linked to target genes annotated as known breast cancer-related drivers in the OncoKB and IntOGen databases [30, 40]. Driver genes with at least 13 enhancer-associated mutations (approximately the top quartile by burden) were selected for downstream analysis.

An oncoplot-like matrix (Fig. 8A) was constructed to visualize the relationships between TF families and the most recurrently mutated breast cancer driver genes. This matrix provides a powerful summary, with each cell representing the proportion of GOF (red) versus LOF (blue) effects on a given TF family for mutations linked to a specific breast cancer-related driver gene. Statistical significance of GOF/LOF enrichment for each relevant gene was assessed using Fisher’s exact test by comparing observed counts against a null distribution generated from 100 trinucleotide-matched simulations via SigProfilerSimulator.

**Fig. 8.**
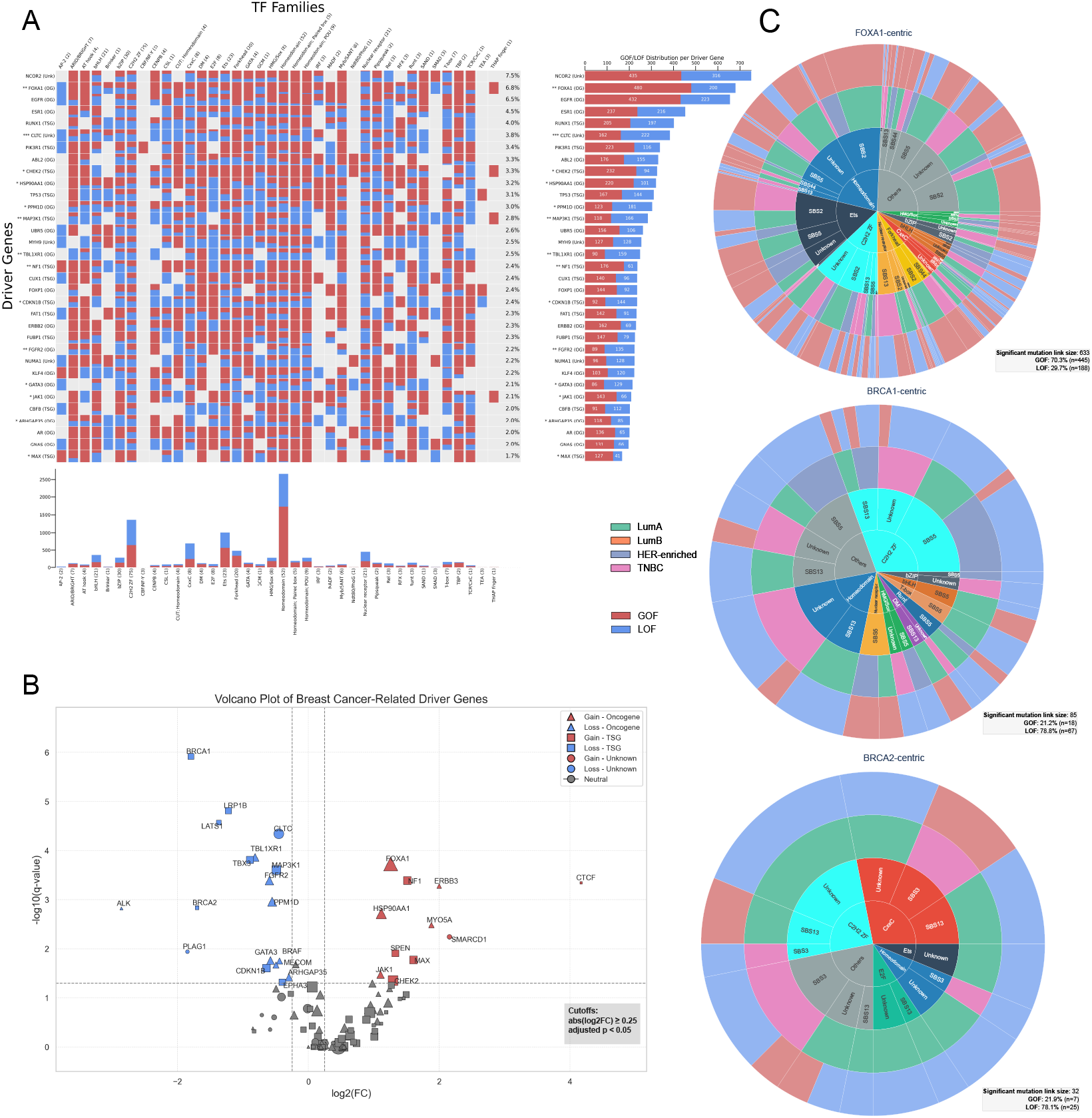
Integration of Enhancer–Target Gene Maps. **A** Oncoplot-like matrix showing TF family–level GOF/LOF proportions for enhancer-linked mutations across top breast cancer driver genes. **B** Volcano plot of driver genes showing GOF/LOF enrichment significance, with markers indicating gene class (OG, TSG, Unknown). **C** Sunburst visualization for *FOXA1, BRCA1* and *BRCA2*, showing enriched GOF and LOF-associated TF family perturbations at their distal enhancer elements.

One prominent example of an enriched driver gene association is *FOXA1*, a known oncogene (OG) in estrogen receptor-positive breast cancer [41]. *FOXA1* target gene in the oncoplot-like matrix had a total of 480 gain-of-function TF-binding mutation events mapped to its enhancer regions, vastly exceeding the random expectation (approximately 120 such events on average in a simulated null background). This corresponds to a significant over-enrichment of mutations that create *de novo* TF binding sites at the *FOXA1* regulatory locus (Fisher’s test, adjusted *p* = 1.86 *×* 10^−4^).

A global volcano plot (Fig. 8B) summarizing the overall distal regulatory perturbation across all targeted driver genes distinguishes between oncogenes (OGs) and tumor suppressor genes (TSGs). Consistent with this pattern, enhancer mutations linked to *FOXA1* preferentially increased binding of Homeodomain, Forkhead, and Ets family TFs (Fig. 8C), whereas canonical tumor suppressors such as *BRCA1* and *BRCA2* were significantly enriched for LOF events, predominantly affecting C2H2-ZF and Homeodomain TF families.

### 3.6 A Multi-scale View of Signature-Driven Effects on Regulatory Gene Programs

We conducted a systematic gene set enrichment analysis to elucidate the higher-order functional consequences of these regulatory perturbations. By linking each predicted TF binding gain or loss prediction to the sets of ABC-predicted target genes and performing enrichment against 14 curated gene programs from the Molecular Signatures Database (MSigDB) Hallmarks and Reactome pathway collections (e.g., DNA Repair, p53 Pathway, E2F Targets, MYC Targets v1/v2, Cell Cycle and Apoptosis), we found that signature-driven TF perturbations converge on non-random downstream gene regulatory programs. The multi-layer structure of these relationships is depicted in Sankey diagrams (Fig. 9, Fig. 10, Fig. 11), which trace flows from BC subtypes to perturbed TF families, onward to enriched target gene programs stratified by direction of regulatory effect (GOF or LOF).

**Fig. 9.**
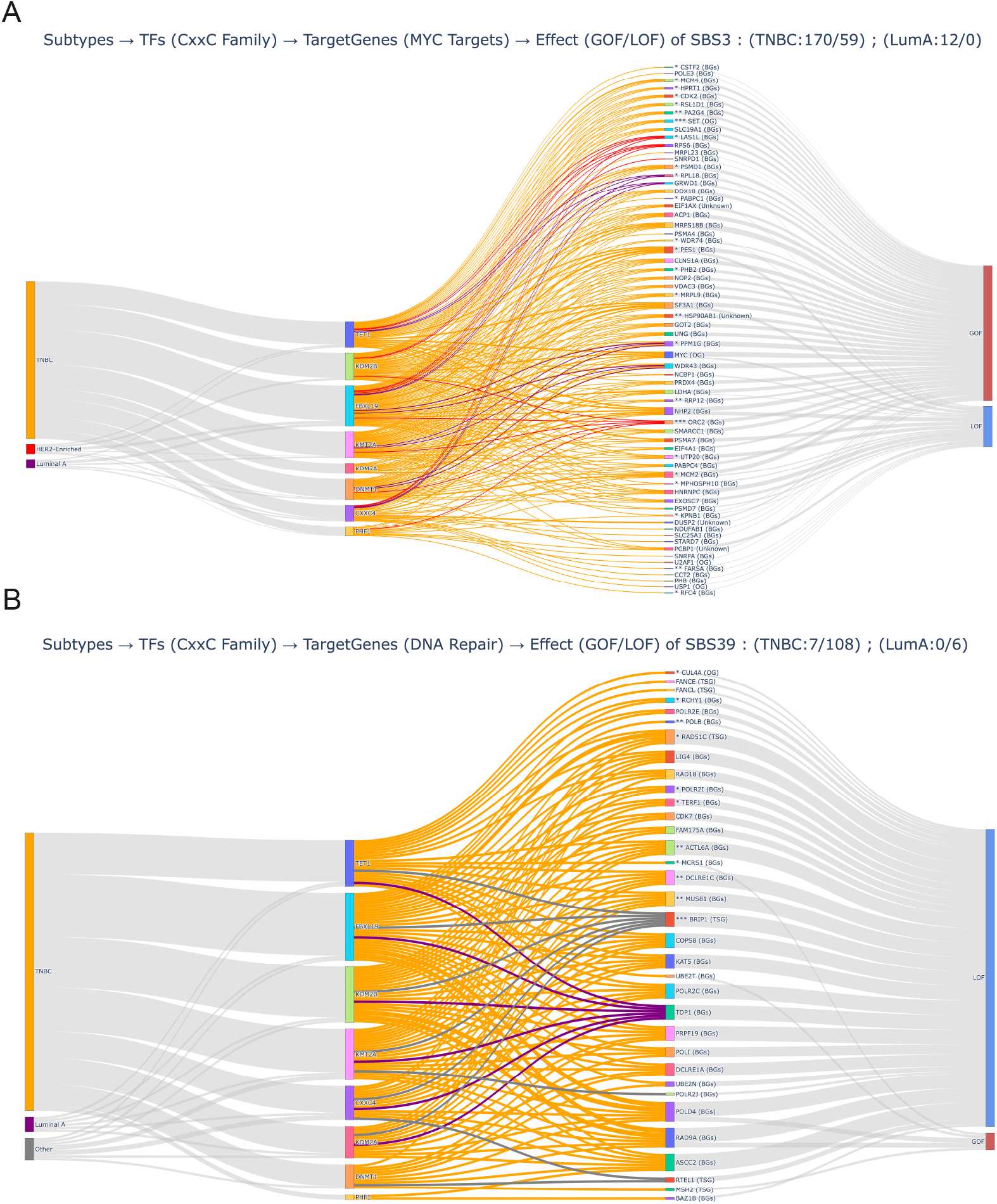
Sankey diagrams illustrating the convergence of TF-family perturbations on downstream gene regulatory programs. Flow width reflects the frequency of identified events, with color coding by breast cancer subtype (TNBC: orange; Luminal A: purple; Luminal B: green; HER2-enriched: red; Other/Not assigned: gray). Statistically significant enrichments are indicated with an asterisk (e.g., *q <* 0.05, based on 100 simulated somatic mutation landscapes from 560 breast cancer samples). **A** SBS3-associated gains of CxxC TF-family binding in TNBC, enriched at enhancers regulating hall-mark MYC target genes. **B** SBS39-associated losses of CxxC TF-family binding in TNBC, enriched at enhancers linked to the *DNA Repair* gene program.

**Fig. 10.**
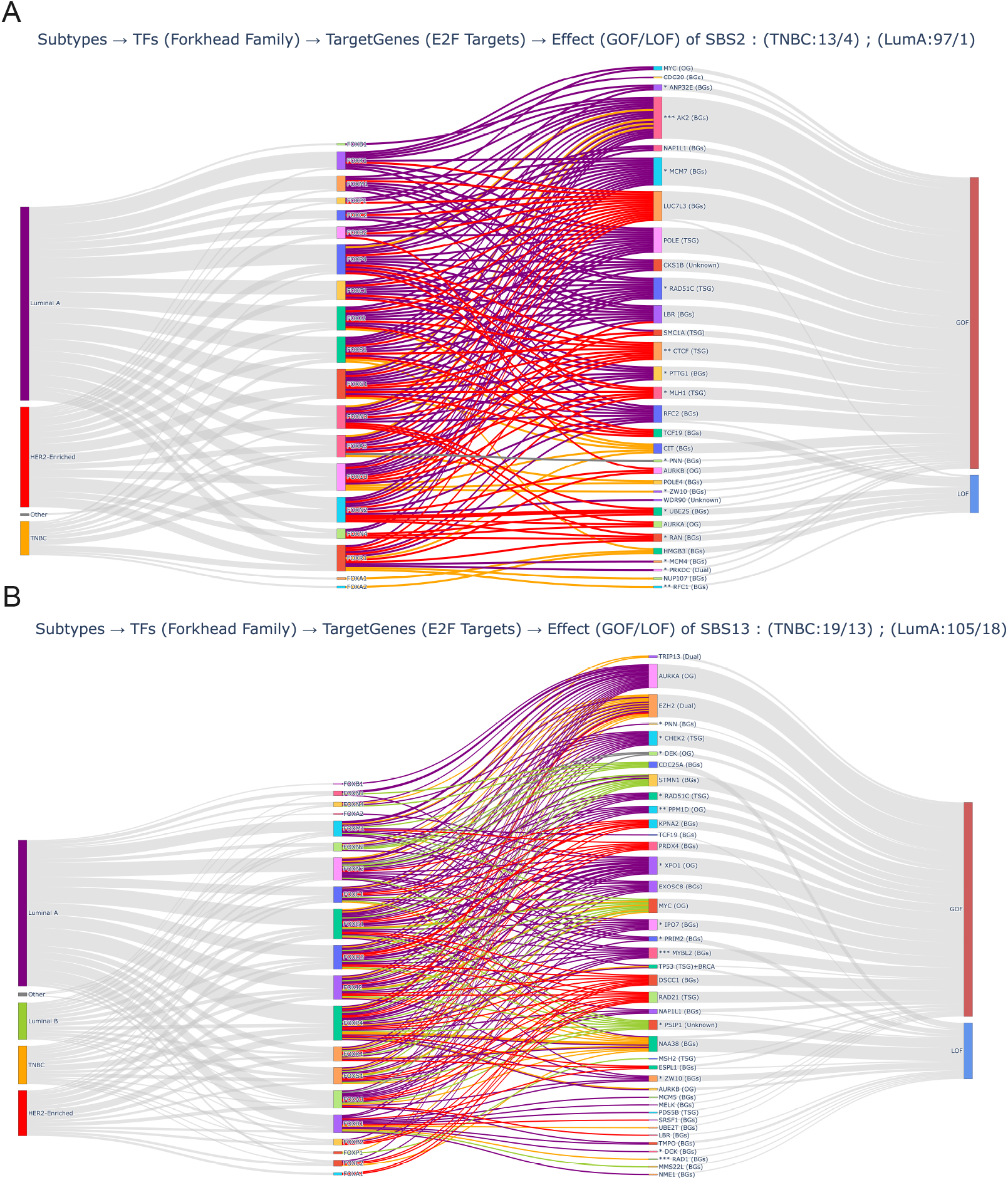
Sankey diagrams illustrating the convergence of TF-family perturbations on downstream gene regulatory programs. Flow width reflects the frequency of identified events; flows are color-coded by breast cancer subtype, and statistically significant enrichments are marked with an asterisk. **A** SBS2 pan-subtype mutations generating gain-of-function binding events in Forkhead TFs, enriched at enhancers regulating E2F target genes (upregulated). **B** SBS13 pan-subtype mutations generating similar Forkhead TF gains, likewise enriched for E2F targets.

**Fig. 11.**
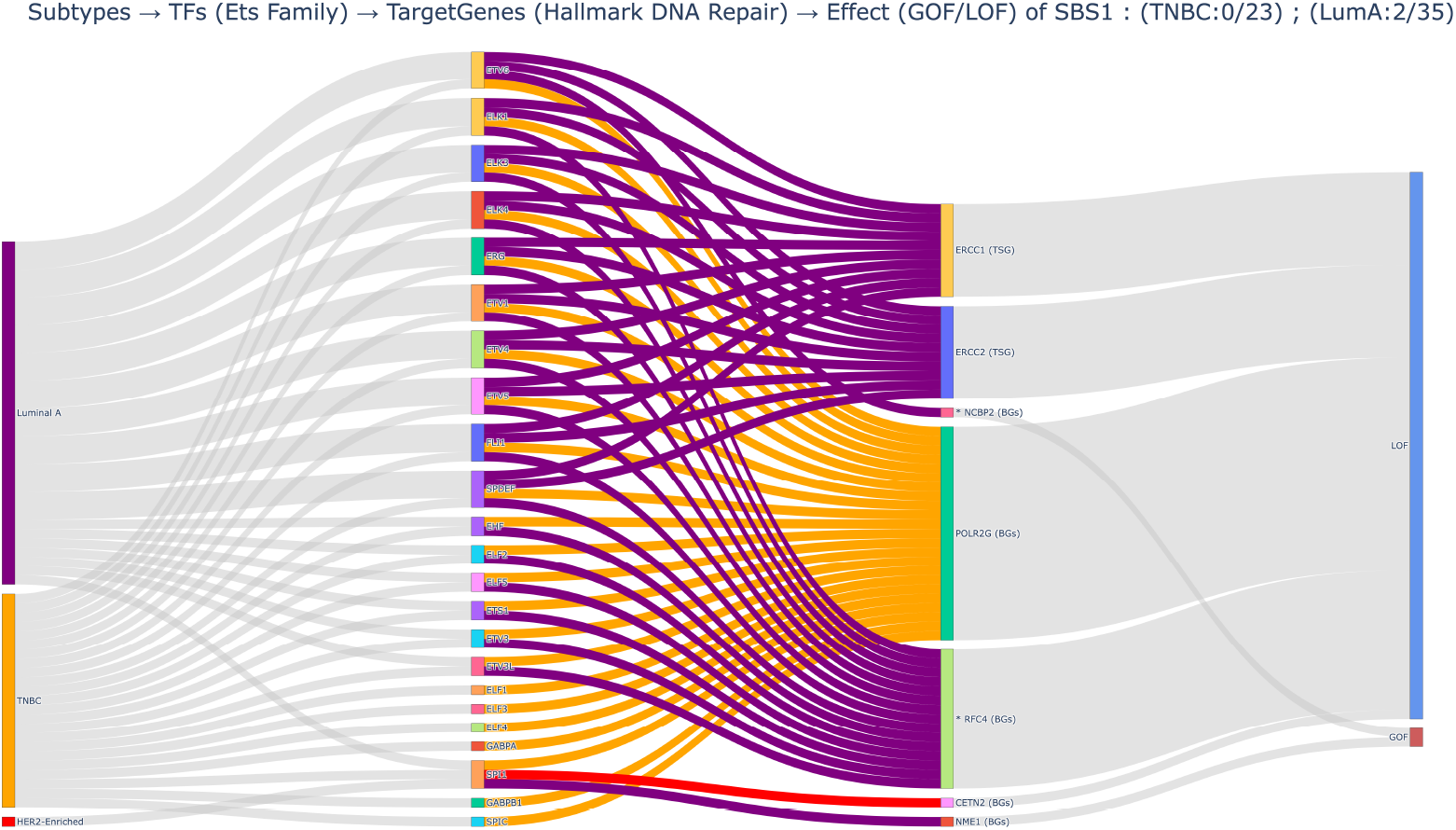
Sankey diagrams for convergence of Ets family perturbations on the regulatory network of genes involved in hallmark DNA repair due to SBS1 mutations.

This analysis specifically revealed that HRD-associated signatures SBS3 and SBS39 within basal-like TNBC samples produced contrasting but mechanistically complementary impacts specifically on CxxC-type zinc-finger TFs (Fig. 9). SBS3-attributed mutations in TNBC samples were found to preferentially create *de novo* binding sites (GOF events) for the CxxC TF family (Fig. 9A). Compared to other luminal subtypes, these GOF perturbations in TNBC were significantly enriched at enhancers regulating the “MYC Targets v1/v2” hallmark gene sets, suggesting a mutational imprint for upregulating MYC-driven proliferative program. In contrast, HRD-associated signature SBS39 predominantly disrupted CxxC TF binding (LOF events) at enhancers linked to the “DNA Repair” gene program within the same TNBC samples (Fig. 9B). Together, these dual and signature-specific effects outline distinct regulatory consequences of HRD-associated mutational processes operating within TNBC.

In addition to subtype-specific effects, we also observed pan-subtype regulatory impacts on gene programs. In our previous predictions, we demonstrated that the APOBEC signatures SBS2 and SBS13 mutations preferentially generated *de novo* binding sites for FOX family TF members (Fig. 6). Enrichment analysis (Fig. 10A, B) revealed that the ABC target genes linked to these gained FOX binding events were highly enriched for E2F target gene sets. E2F target genes play a central role in controlling cell-cycle progression, particularly the G1/S transition, and their dysregulation is a hallmark of uncontrolled proliferation in breast cancer. Our findings align with prior observations that E2F-regulated genes are commonly overexpressed across diverse BC subtypes [42]. The flows from SBS2/SBS13 to FOX members and onward to E2F target genes were among the most significant GOF patterns observed (asterisks in Fig. 10) compared to the null distribution.

Conversely, the age-related signature SBS1, previously associated with a loss of Ets family bindings (Fig. 7), was found to drive LOF events at enhancers regulating DNA Repair genes (Fig. 11). Within this context, the *POLR2G* gene emerged as a notable target, with multiple Ets family members showing SBS1-associated LOF perturbations originating exclusively from TNBC samples. Other Ets-linked enhancer–gene associations were more frequently observed in Luminal A tumors, highlighting subtype-specific divergence in SBS1-driven regulatory effects.

## 4 Discussion

Despite the growing recognition of non-coding mutations in cancer biology, the systematic assessment of how mutational signatures perturb TF binding and downstream regulatory programs remain underexplored. Specifically, few studies have integrated mutational signature analysis with predicted TF binding alterations [36, 37]. We address this gap by applying a comprehensive *in silico* framework to 560 breast cancer genomes within a subtype-specific context. Our approach integrates mutational signatures, perturbed TF binding landscapes, and enhancer–gene linkages to infer gene regulatory consequences. For this purpose, we built 6-mer-based regression models with SGD for 403 human TFs that achieved moderate goodness of fit (mean *R*^2^ ≈ 0.39, median *R*^2^ ≈ 0.35). The variability in our models likely reflects noisy differences in the nature of the training data (ChIP-seq vs. PBM), but overall, the model panel remained sufficiently robust to detect meaningful TF-DNA affinity changes. By scanning ≈ 3.5 million single-base substitutions from the cohort, we identified statistically significant cases of TF binding affinity changes, such as the *de novo* creation of an MYBL2 motif or the disruption of an ETS1 site, that align with known binding specificities (Fig. 3). These results demonstrate the utility of our approach in prioritizing candidate regulatory mutations from a vast mutational background.

Different mutational signatures leave distinct “fingerprints” on TF binding landscapes by preferentially affecting specific TFs in a directional manner. For instance, APOBEC signatures (SBS2 and SBS13) and the HRD signature SBS3 primarily produce gain-of-function events in FOX and CxxC sites, respectively. Conversely, clock-like SBS1 and HRD-associated SBS39 lead to loss-of-function events in ETS and CxxC sites, respectively. These signature-specific perturbations suggest that mutational mechanisms do not act randomly, but rather shape transcriptional networks by systematically altering TF binding patterns.

Integrating predicted TF binding changes with ABC-derived enhancer–target gene maps revealed that these perturbations converge on critical cancer-related gene programs rather than occurring randomly. For instance, GOF mutations frequently cluster at enhancers of proliferation-associated genes (such as MYC transcriptional targets or E2F-regulated cell cycle genes), whereas LOF mutations tend to affect enhancers of DNA repair genes. These patterns suggest the regulatory genome is actively sculpted during tumorigenesis to reinforce cancer hallmarks, effectively by “pressing the accelerator” on growth-promoting pathways while “cutting the brakes” on genomic maintenance and tumor suppression.

We observed that the regulatory impact of mutational signatures is modulated by breast cancer molecular subtypes. Consistent with studies linking HRD-associated signatures to TNBC pathogenesis [32, 34], we found that CXXC family TF enrichment—GOF under SBS3 and LOF under SBS39—is largely driven by basal-like TNBC samples. Specifically, SBS3 mutations preferentially target enhancers of MYC-regulated genes, potentially promoting proliferative signaling. This matches reports of MYC hyperactivation, which is linked to proliferation and immune suppression in aggressive TNBC [43, 44]. Moreover, the CXXC-family regulator KDM2B is known to cooperate with MYC to drive oncogenic transcription [45]. This supports our finding that SBS3-induced CXXC gain motifs are enriched at MYC target enhancers. More broadly, our modeling suggests a framework where distinct mutational signatures preferentially perturb specific transcriptional networks.

Despite the insights gained, our study has several limitations. First, our findings rely on *in silico* predictions that require experimental validation. While we infer TF binding alterations from sequence-based modeling and association analyses, future studies utilizing allele-specific ChIP-seq or CRISPR-mediated base editing of enhancers are essential. These steps will confirm whether predicted mutations functionally alter binding and modulate gene expression *in vivo*. Second, our models do not fully capture cell-type-specific chromatin context. Actual TF binding in tumors is governed by chromatin accessibility and specific histone modifications. Our sequence-based approach cannot account for cases where a strong motif might be inaccessible or, conversely, a weak motif is rendered functional by the local chromatin environment. While we partially mitigated this by restricting our analysis to putative regulatory regions via ABC enhancer maps, these models are derived from generalized datasets. Consequently, they may not fully capture the patient-specific chromatin architecture or the unique epigenetic remodeling characteristic of individual tumors. Third, our framework incorporates inherent biases from external data sources, such as the assumed generalizability of PBM or ChIP-seq-derived motifs and the accuracy of ABC-based enhancer-gene linkages. Variability in TF model performance, particularly for factors characterized by low-quality ChIP-seq data, may lead to underestimation of binding alterations for certain transcription factors.

Nonetheless, this framework carries broad implications for computational oncology. The preferential targeting of specific gene networks by distinct mutational signatures reveals a novel interaction between genomic instability and gene regulatory circuitry. Practically, our approach offers a scalable means to prioritize non-coding driver candidates by narrowing the vast mutation space to high-priority, functionally interpretable sites. Consistent with recent findings that TF occupancy is driven by the aggregate effect of multiple overlapping binding sites rather than by a single high-affinity motif [46], our model does not treat mutations in isolation; instead, it quantifies the cumulative impact on all overlapping 6-mer features within an 11-bp window, providing a robust *k* -mer-based approximation of the local binding architecture.

However, our *k* -mer regression model is inherently limited by its 6-mer feature space and 11-bp window. Consequently, very weak binding sites, which often rely on long-range interactions or complex sequence contexts, may be underrepresented. Our estimates of mutational impact are therefore likely conservative; future refinements could incorporate more expressive architectures, such as convolutional neural networks or transformers, to capture dependencies beyond the current 11-bp limit. Moving forward, experimental validation of these *in silico* hypotheses remains a critical next step. Expanding this framework to pan-cancer datasets will further clarify whether the observed signature-TF-pathway linkages are unique to breast cancer or represent universal principles of regulatory disruption. Future research should prioritize the integration of transcriptomic profiles with tissue-specific epigenomic data, such as single-cell ATAC-seq, to bridge the gap between predicted binding perturbations and downstream changes in gene expression. Finally, as this study focused on single-base substitutions, incorporating other mutational events such as doublet-base substitutions, indels and structural variants will offer a more comprehensive view of the regulatory landscape.

## 5 Conclusion

This study investigates the impact of somatic mutations on TF binding and downstream regulatory programs across 560 breast cancer samples, characterizing how these alterations are driven by specific mutational processes and varied across molecular subtypes. Using a robust *in silico* pipeline, we quantitatively predicted the impact of approximately 3.5 million somatic mutations on TF binding affinity, uncovering distinct signature-specific and subtype-specific trends. Our analysis reveals that mutational signatures attributed to APOBEC cytidine deamination, age-related methylation damage, and HRD-associated processes induce distinct patterns of TF family perturbations and gene regulatory outcomes. These findings reveal a novel connection between a tumor’s mutational profile and its gene regulatory network.

In conclusion, our work establishes a robust foundation for systematically linking mutational processes to their downstream gene regulatory consequences. This scalable framework enables the prioritization of functional non-coding mutations by characterizing both gain- and loss-of-function effects on TF binding within the vast landscapes of cancer genomes. Ultimately, these findings underscore the importance of interpreting somatic alterations through the lens of their underlying mutational processes.

## Supplementary information

Supplementary information is available for this paper and includes additional figures (Figs. S1–S4), providing further methodological details and extended results.

## Acknowledgements

Not applicable.

## Declarations

### Funding

This research received no specific grant from any funding agency in the public, commercial, or not-for-profit sectors.

### Competing interests

The authors declare that they have no competing interests.

### Ethics approval and consent to participate

Not applicable.

### Consent for publication

Not applicable.

### Availability of data and materials

The datasets generated and/or analyzed during the current study are available from the corresponding author on reasonable request.

### Code availability

The source code used in this study is available from the corresponding author on reasonable request.

### Authors’ contributions

HHK developed the computational framework and drafted the manuscript. BO supervised the study and contributed to manuscript revision. All authors read and approved the final manuscript.

## Notes

### Competing Interest Statement

The authors have declared no competing interest.

## References

[1] Khurana E, Fu Y, Chakravarty D, Demichelis F, Rubin MA, Gerstein M. Role of non-coding sequence variants in cancer. Nature Reviews Genetics 2016 17:2. 2016 Jan;17(2):93–108. Publisher: Nature Publishing Group. 10.1038/nrg.2015.17.

[2] Maurano MT, Humbert R, Rynes E, Thurman RE, Haugen E, Wang H, et al. Systematic localization of common disease-associated variation in regulatory DNA. Science (New York, NY). 2012 Sep;337(6099):1190–1195. Publisher: Science. 10.1126/SCIENCE.1222794.

[3] Johnson DS, Mortazavi A, Myers RM, Wold B. Genome-wide mapping of in vivo protein-DNA interactions. Science (New York, NY). 2007 Jun;316(5830):1497– 1502. 10.1126/science.1141319.

[4] Jolma A, Yan J, Whitington T, Toivonen J, Nitta KR, Rastas P, et al. DNA-binding specificities of human transcription factors. Cell. 2013 Jan;152(1-2):327– 339. 10.1016/j.cell.2012.12.009.

[5] Rockel S, Geertz M, Maerkl SJ. MITOMI: a microfluidic platform for in vitro characterization of transcription factor-DNA interaction. Methods in Molecular Biology (Clifton, NJ). 2012;786:97–114. 10.1007/978-1-61779-292-26.

[6] Berger MF, Philippakis AA, Qureshi AM, He FS, Estep PW, Bulyk ML. Compact, universal DNA microarrays to comprehensively determine transcription-factor binding site specificities. Nature Biotechnology. 2006 Nov;24(11):1429– 1435. Publisher: Nature Publishing Group. 10.1038/nbt1246.

[7] Fostier J. BLAMM: BLAS-based algorithm for finding position weight matrix occurrences in DNA sequences on CPUs and GPUs. BMC Bioinformatics. 2020 Mar;21(2):1–13. Publisher: BioMed Central Ltd.. 10.1186/S12859-020-3348-6/FIGURES/4.

[8] Zia A, Moses AM. Towards a theoretical understanding of false positives in DNA motif finding. BMC Bioinformatics. 2012 Jun;13(1):1–9. Publisher: BioMed Central. 10.1186/1471-2105-13-151/FIGURES/6.

[9] Bi Y, Kim H, Gupta R, Davuluri RV. Tree-Based Position Weight Matrix Approach to Model Transcription Factor Binding Site Profiles. PLoS ONE. 2011 Sep;6(9):e24210. 10.1371/JOURNAL.PONE.0024210.

[10] Bulyk ML, Johnson PLF, Church GM. Nucleotides of transcription factor binding sites exert interdependent effects on the binding affinities of transcription factors. Nucleic Acids Research. 2002 Mar;30(5):1255–1261. 10.1093/nar/30.5.1255.

[11] Pique-Regi R, Degner JF, Pai AA, Gaffney DJ, Gilad Y, Pritchard JK. Accurate inference of transcription factor binding from DNA sequence and chromatin accessibility data. Genome Research. 2011 Mar;21(3):447–455. Company: Cold Spring Harbor Laboratory Press Distributor: Cold Spring Harbor Laboratory Press Institution: Cold Spring Harbor Laboratory Press Label: Cold Spring Harbor Laboratory Press Publisher: Cold Spring Harbor Lab. 10.1101/gr.112623.110.

[12] Behjati Ardakani F, Schmidt F, Schulz MH. Predicting transcription factor binding using ensemble random forest models. F1000Research. 2019 Sep;7:1603. 10.12688/f1000research.16200.2.

[13] Martin V, Zhao J, Afek A, Mielko Z, Gordân R. QBiC-Pred: quantitative predictions of transcription factor binding changes due to sequence variants. Nucleic Acids Research. 2019 Jul;47(W1):W127–W135. 10.1093/nar/gkz363.

[14] Alipanahi B, Delong A, Weirauch MT, Frey BJ. Predicting the sequence specificities of DNA- and RNA-binding proteins by deep learning. Nature Biotechnology. 2015 Aug;33(8):831–838. 10.1038/nbt.3300.

[15] Quang D, Xie X. FactorNet: A deep learning framework for predicting cell type specific transcription factor binding from nucleotide-resolution sequential data. Methods. 2019 Aug;166:40–47. 10.1016/j.ymeth.2019.03.020.

[16] Wang K, Zeng X, Zhou J, Liu F, Luan X, Wang X. BERT-TFBS: a novel BERT-based model for predicting transcription factor binding sites by transfer learning. Briefings in Bioinformatics. 2024 May;25(3):bbae195. 10.1093/bib/bbae195.

[17] Gao Z, Liu Q, Zeng W, Jiang R, Wong WH. EpiGePT: a pretrained transformer-based language model for context-specific human epigenomics. Genome Biology. 2024 Dec;25(1):310. 10.1186/s13059-024-03449-7.

[18] Alexandrov LB, Nik-Zainal S, Wedge DC, Aparicio SAJR, Behjati S, Biankin AV, et al. Signatures of mutational processes in human cancer. Nature. 2013;500(7463):415–421. Publisher: Nature Publishing Group. 10.1038/nature12477.

[19] Steinhaus R, Robinson PN, Seelow D. FABIAN-variant: predicting the effects of DNA variants on transcription factor binding. Nucleic Acids Research. 2022 Jul;50(W1):W322–W329. 10.1093/nar/gkac393.

[20] Zhao J, Li D, Seo J, Allen AS, Gordân R. Quantifying the Impact of Non-coding Variants on Transcription Factor-DNA Binding. In: Sahinalp SC, editor. Research in Computational Molecular Biology. Cham: Springer International Publishing; 2017. p. 336–352.

[21] Wilkinson L, Gathani T. Understanding breast cancer as a global health concern. The British Journal of Radiology. 2022 Feb;95(1130):20211033. 10.1259/bjr.20211033.

[22] Nik-Zainal S, Davies H, Staaf J, Ramakrishna M, Glodzik D, Zou X, et al. Landscape of somatic mutations in 560 breast cancer whole genome sequences. Nature. 2016 May;534(7605):47–54. 10.1038/nature17676.

[23] Nasser J, Bergman DT, Fulco CP, Guckelberger P, Doughty BR, Patwardhan TA, et al. Genome-wide enhancer maps link risk variants to disease genes. Nature. 2021 May;593(7858):238–243. Publisher: Nature Publishing Group. 10.1038/s41586-021-03446-x.

[24] Dunham I, Kundaje A, Aldred SF, Collins PJ, Davis CA, Doyle F, et al. An integrated encyclopedia of DNA elements in the human genome. Nature. 2012 Sep;489(7414):57–74. Publisher: Nature Publishing Group. 10.1038/nature11247.

[25] Hume MA, Barrera LA, Gisselbrecht SS, Bulyk ML. UniPROBE, update 2015: new tools and content for the online database of protein-binding microarray data on protein–DNA interactions. Nucleic Acids Research. 2015 Jan;43(D1):D117–D122. 10.1093/nar/gku1045.

[26] Weirauch M, Yang A, Albu M, Cote AG, Montenegro-Montero A, Drewe P, et al. Determination and Inference of Eukaryotic Transcription Factor Sequence Specificity. Cell. 2014 Sep;158(6):1431–1443. 10.1016/j.cell.2014.08.009.

[27] Islam SMA, Díaz-Gay M, Wu Y, Barnes M, Vangara R, Bergstrom EN, et al. Uncovering novel mutational signatures by de novo extraction with SigProfilerExtractor. Cell Genomics. 2022 Nov;2(11). Publisher: Elsevier. 10.1016/j.xgen.2022.100179.

[28] Sondka Z, Dhir NB, Carvalho-Silva D, Jupe S, Madhumita, McLaren K, et al. COSMIC: a curated database of somatic variants and clinical data for cancer. Nucleic Acids Research. 2024 Jan;52(D1):D1210–D1217. 10.1093/nar/gkad986.

[29] Bergstrom EN, Barnes M, Martincorena I, Alexandrov LB.: Generating realistic null hypothesis of cancer mutational landscapes using SigProfilerSimulator. bioRxiv. Pages: 2020.02.13.948422 Section: New Results. Available from: https://www.biorxiv.org/content/10.1101/2020.02.13.948422v1.

[30] Chakravarty D, Gao J, Phillips S, Kundra R, Zhang H, Wang J, et al. OncoKB: A Precision Oncology Knowledge Base. JCO Precision Oncology. 2017 May;(1):1– 16. Publisher: Wolters Kluwer. 10.1200/PO.17.00011.

[31] Pratt HE, Andrews GR, Phalke N, Huey JD, Purcaro M, van der Velde A, et al. Factorbook: an updated catalog of transcription factor motifs and candidate regulatory motif sites. Nucleic Acids Research. 2022 Jan;50(D1):D141–D149. 10.1093/nar/gkab1039.

[32] Pan JW, Tan ZC, Ng PS, Zabidi MMA, Nur Fatin P, Teo JY, et al. Gene expression signature for predicting homologous recombination deficiency in triplenegative breast cancer. npj Breast Cancer. 2024 Jul;10(1):60. Publisher: Nature Publishing Group. 10.1038/s41523-024-00671-1.

[33] Yao S, Wei L, Hu Q, Liu S, Manojlovic Z, Fiorica PN, et al. Mutational landscape of triple-negative breast cancer in African American women. Nature Genetics. 2025 Sep;57(9):2166–2176. Publisher: Nature Publishing Group. 10.1038/s41588-025-02322-y.

[34] Shah B, Hussain M, Seth A. Homologous Recombination Deficiency in Ovarian and Breast Cancers: Biomarkers, Diagnosis, and Treatment. Current Issues in Molecular Biology. 2025 Aug;47(8):638. Publisher: Multidisciplinary Digital Publishing Institute. 10.3390/cimb47080638.

[35] Dananberg A, Striepen J, Rozowsky JS, Petljak M. APOBEC Mutagenesis in Cancer Development and Susceptibility. Cancers. 2024 Jan;16(2):374. Number: 2 Publisher: Multidisciplinary Digital Publishing Institute. 10.3390/cancers16020374.

[36] Liu M, Boot A, Ng AWT, Gordân R, Rozen SG. Mutational processes in cancer preferentially affect binding of particular transcription factors. Scientific Reports. 2021 Feb;11(1):3339. Publisher: Nature Publishing Group. 10.1038/s41598-021-82910-0.

[37] Yiu Chan CW, Gu Z, Bieg M, Eils R, Herrmann C. Impact of cancer mutational signatures on transcription factor motifs in the human genome. BMC Medical Genomics. 2019 May;12(1):64. 10.1186/s12920-019-0525-4.

[38] Li Z, Abraham BJ, Berezovskaya A, Farah N, Liu Y, Leon T, et al. APOBEC signature mutation generates an oncogenic enhancer that drives LMO1 expression in T-ALL. Leukemia. 2017 Oct;31(10):2057–2064. Publisher: Nature Publishing Group. 10.1038/leu.2017.75.

[39] Spisak N, de Manuel M, Milligan W, Sella G, Przeworski M. Disentangling sources of clock-like mutations in germline and soma. bioRxiv. 2023 Sep;p. 2023.09.07.556720. 10.1101/2023.09.07.556720.

[40] Martínez-Jiménez F, Muiños F, Sentís I, Deu-Pons J, Reyes-Salazar I, Arnedo-Pac C, et al. A compendium of mutational cancer driver genes. Nature Reviews Cancer. 2020 Oct;20(10):555–572. Publisher: Nature Publishing Group. 10.1038/s41568-020-0290-x.

[41] Seachrist DD, Anstine LJ, Keri RA. FOXA1: A Pioneer of Nuclear Receptor Action in Breast Cancer. Cancers. 2021 Jan;13(20):5205. Number: 20 Publisher: Multidisciplinary Digital Publishing Institute. 10.3390/cancers13205205.

[42] Nikonezhad B, Lotfian M, Manavi N, Zamani A, Mahdevar M. Insights into the E2F target genes in breast cancer: associations of pathway genes with prognosis and immune cell filtration based on in silico and ex vivo analyses. Cancer Cell International. 2025 Jun;25:203. 10.1186/s12935-025-03839-2.

[43] Schulze A, Oshi M, Endo I, Takabe K. MYC Targets Scores Are Associated with Cancer Aggressiveness and Poor Survival in ER-Positive Primary and Metastatic Breast Cancer. International Journal of Molecular Sciences. 2020 Oct;21(21):8127. 10.3390/ijms21218127.

[44] Zimmerli D, Brambillasca CS, Talens F, Bhin J, Linstra R, Romanens L, et al. MYC promotes immune-suppression in triple-negative breast cancer via inhibition of interferon signaling. Nature Communications. 2022 Nov;13(1):6579. Publisher: Nature Publishing Group. 10.1038/s41467-022-34000-6.

[45] Chavdoula E, Anastas V, La Ferlita A, Aldana J, Carota G, Spampinato M, et al. Transcriptional regulation of amino acid metabolism by KDM2B, in the context of ncPRC1.1 and in concert with MYC and ATF4. Metabolism: Clinical and Experimental. 2024 Jan;150:155719. 10.1016/j.metabol.2023.155719.

[46] Khetan S, Carroll BS, Bulyk ML. Multiple overlapping binding sites determine transcription factor occupancy. Nature. 2025 Oct;646(8086):1001–1011. 10.1038/s41586-025-09472-3.

