## Supplementary Information for "A Robust Framework for Predicting Mutation Effects on Transcription Factor Binding: Insights from Mutational Signatures in 560 Breast Cancer Genomes"

### 1. INTRODUCTION

This Supplementary Information provides additional analyses, figures, and tables that support and extend the results presented in the main manuscript. In particular, it includes extended performance results, robustness analyses of the proposed sequence-based models, and additional stratifications by mutational signatures, transcription factor families, and subtype-specific analyses that could not be fully accommodated in the main text due to space constraints. All supplementary figures and tables are referenced in the main manuscript and are intended to facilitate transparency, reproducibility, and deeper inspection of the computational framework.

### 2. SUPPLEMENTARY FIGURES

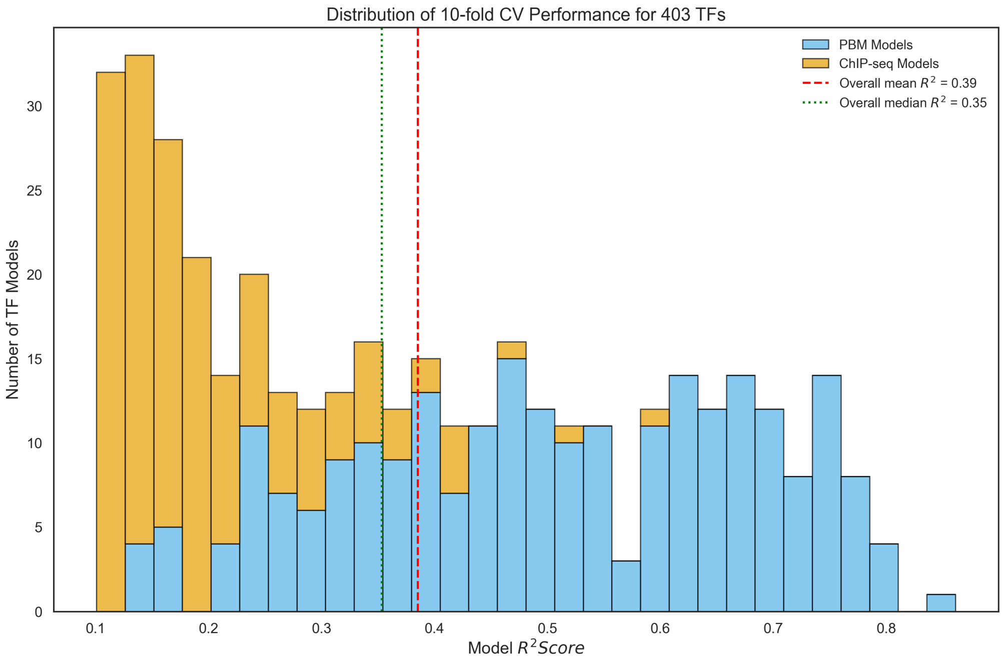


**Fig. S1.** Distribution of predictive performance across transcription factor (TF) binding models. Histogram showing the distribution of 10-fold cross-validated model performance ($R^2$) for the final set of 403 transcription factor (TF) binding models. Models trained on in vitro protein-binding microarray (PBM) data are shown in blue, while models trained on in vivo ChIP-seq data are shown in orange. The dashed red line indicates the overall mean $R^2$ (0.39), and the dotted green line denotes the overall median $R^2$ (0.35). Consistent with differences in data quality, PBM-based models generally achieve higher predictive performance than ChIP-seq-based models, motivating the use of tiered model inclusion thresholds ($R^2 > 0.15$ for PBM models and $R^2 > 0.10$ for ChIP-seq models).

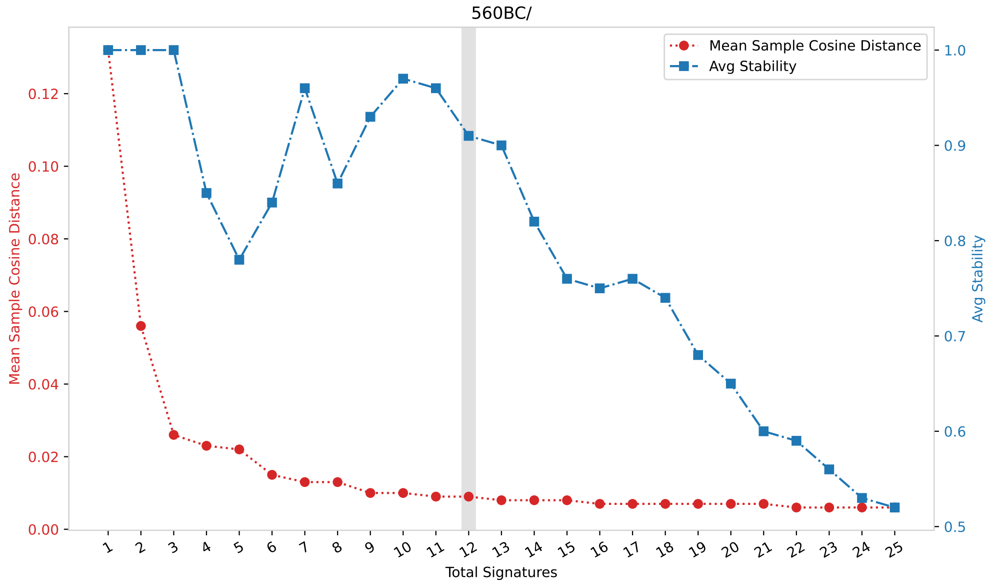


**Fig. S2.** Selection of the optimal number of de novo single-base substitution (SBS) signatures. Signature selection plot obtained from de novo SBS signature extraction using SigProfilerExtractor across the 560 breast cancer genomes. The mean sample cosine distance (red, left y-axis) reflects reconstruction error between the original and reconstructed mutational profiles, while average signature stability (blue, right y-axis) quantifies the reproducibility of extracted signatures across repeated runs. The shaded vertical region indicates the selected solution of 12 SBS signatures, which represents a balance between low reconstruction error and high stability. These 12 de novo signatures were subsequently used for downstream analyses and decomposed into COSMIC v3.4 reference signatures.

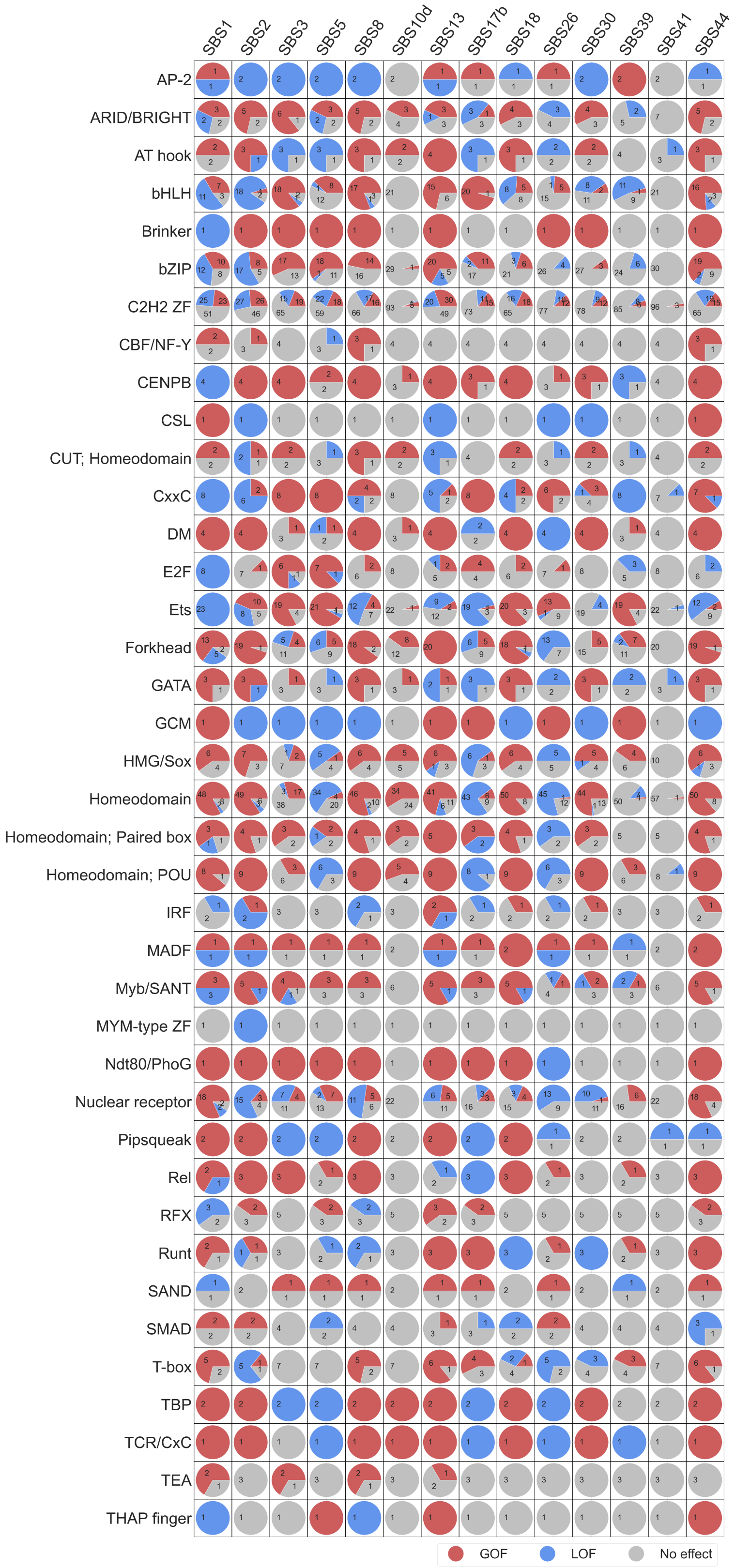


**Fig. S3.** (See legend on next page.)

(See figure on previous page.)

**Fig S3. Family-level aggregation of transcription factor binding perturbations across mutational signatures.** Heatmap of pie charts summarizing transcription factor (TF) family–level binding perturbations across mutational signatures in the 560 breast cancer genomes. Rows correspond to TF families grouped by shared DNA-binding domain (DBD) architecture, and columns represent single-base substitution (SBS) mutational signatures. For each signature–family pair, pie chart segments indicate the proportion of TF family members exhibiting significant gain-of-function (GOF; red), loss-of-function (LOF; blue), or non-significant (gray) predicted binding effects. This family-level aggregation highlights coordinated and non-random patterns of regulatory disruption, with several mutational signatures showing pronounced directional biases toward GOF or LOF effects within specific TF families.

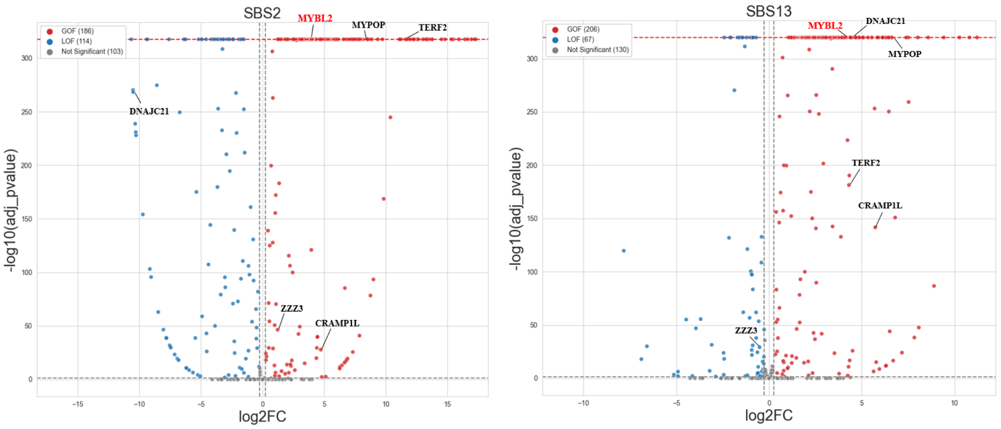


**A** **B**

**Fig. S4.** APOBEC-associated gain-of-function enrichment of Myb/SANT family transcription factors. Volcano plots summarizing the differential effects of APOBEC-associated mutational signatures on transcription factor (TF) binding for SBS2 **(A)** and SBS13 **(B)**. The x-axis shows the log2 fold change (log2FC) of predicted TF binding effects, while the y-axis represents statistical significance (-log10 adjusted p-value). TFs exhibiting significant gain-of-function (GOF) effects are shown in red, loss-of-function (LOF) effects in blue, and non-significant TFs in gray. Dashed vertical lines indicate the effect size thresholds used to classify GOF and LOF events. Across both SBS2 and SBS13, members of the Myb/SANT family display a pronounced shift toward GOF effects, consistent with preferential enhancement of MYB-like binding motifs by APOBEC-driven mutations. Selected representative TFs are highlighted.
